# Pyochelin biotransformation by *Staphylococcus aureus* shapes bacterial competition with *Pseudomonas aeruginosa* in polymicrobial infections

**DOI:** 10.1101/2022.04.18.486787

**Authors:** C. Jenul, K. Keim, J. Jens, M. J. Zeiler, K. Schilcher, M. Schurr, C. Melander, V.V. Phelan, A.R. Horswill

**Author notes:** Corresponding author. (ARH) (VVP).

## Abstract

*Pseudomonas aeruginosa* and *Staphylococcus aureus* are among the most frequently isolated bacterial species from polymicrobial infections of cystic fibrosis patients and chronic wounds. We applied mass spectrometry guided interaction studies to determine how chemical interaction shapes the fitness and community structure during co-infection of these two pathogens. We demonstrate that *S. aureus* is equipped with an elegant mechanism to inactivate pyochelin via the yet uncharacterized methyltransferase Spm (<u>s</u>taphylococcal <u>p</u>yochelin <u>m</u>ethyltransferase). Methylation of pyochelin abolishes the siderophore activity of pyochelin and significantly lowers pyochelin-mediated intracellular ROS production in *S. aureus*. In a murine wound co-infection model, a *S. aureus* mutant unable to methylate pyochelin shows significantly lower fitness as compared to its parental strain. Thus, Spm mediated pyochelin methylation is a novel mechanism to increase *S. aureus* survival during *in vivo* competition with *P. aeruginosa*.

## Introduction

*Staphylococcus aureus* and *Pseudomonas aeruginosa* are opportunistic pathogens frequently co-isolated from sites of chronic infection, including the pulmonary infections of persons with cystic fibrosis (CF)^1,2^ and chronic wounds^3–8^. In CF pulmonary infections, co-infections with these two pathogens are associated with worsened patient outcomes due to increased inflammation, virulence factor expression, and antibiotic tolerance^1,9,10^. Contrary to CF lung infections, the impact of *S. aureus*/*P. aeruginosa* co-infection in chronic wounds is not well understood. Due to the lack of comprehensive long-term clinical data on chronic wound co-infection, studies have primarily relied on *in vitro*, *ex vivo*, and animal models to better understand how co-infection with *S. aureus* and *P. aeruginosa* influences chronic wound outcomes^11–13^. Despite frequent co-isolation from the infection environment, which suggests some level of co-existence, co-cultivation under standard laboratory conditions leads to antagonistic interactions between these two microbes, with *P. aeruginosa* outcompeting *S. aureus*^12,14–17^. This phenomenon is highly strain-dependent and influenced by the inoculum ratio of the two bacterial species as well as culturing conditions^18^.

*P. aeruginosa* isolates from initial infection of the lungs of people with CF are antagonistic towards *S. aureus*, while isolates from chronic infections may support co-existence^17^. Research into chronic *P. aeruginosa* infection has shown that its prolonged persistence in the CF environment results in the accumulation of mutations, leading to increased biofilm formation and antibiotic resistance, decreased virulence, and general metabolic adaption to the CF environment, which includes a shift from siderophore-based iron acquisition to heme-based iron acquisition^19–22^. Importantly, chronic wound biopsies suggest that *S. aureus* and *P. aeruginosa* occupy distinct niches and, therefore, may interact through chemical competition rather than physical interaction^23^.

*S. aureus* is a known early colonizer of cystic fibrosis patients and the main pathogen found in young children, while *P. aeruginosa* is the dominant pathogen found in older CF patients^24^. In addition, *P. aeruginosa* is not commonly found to inhabit the skin^25^, while *S. aureus* is a common, although transient, member of the skin microbiota of healthy individuals^25^ and is likely seeded from its primary niche, the anterior nares^26^. Taken together, *S. aureus* is likely to encounter non-adapted *P. aeruginosa* strains during the early stages of skin wound infections as well as during the early onset of *P. aeruginosa* CF lung infection. The objective of this study was to determine the roles of key metabolic exchange factors governing the *in vivo* co-existence of non-adapted *P. aeruginosa* and *S. aureus*.

The antagonistic interactions between *S. aureus* and *P. aeruginosa* are multifactorial. *P. aeruginosa* produces a variety of extracellular factors that kill or inhibit growth of *S. aureus*, including the phenazine respiratory toxins, such as pyocyanin^27^; alkyl-hydroxyquinoline *N*-oxides, a class of respiratory toxins known to inhibit *S. aureus* growth^28^; the metalloendopeptidase LasA that lyses *S. aureus* via peptidoglycan cleavage^29,30^; rhamnolipids, which have anti-adhesive effects and disperse staphylococcal biofilms^31,32^; and the siderophores pyoverdine^33,34^ and pyochelin^34,35^, with the latter not only exhibiting iron chelating activity but also inducing intracellular ROS production in *S. aureus* and other bacteria^36,37^. In response, *S. aureus* forms small-colony variants (SCV)^16^, a phenotype that is selected for when *S. aureus* is subjected to respiratory toxins like pyocyanin and alkyl-hydroxyquinoline N-oxides^38,39^.

Herein, we identify an additional mechanism that *S. aureus* uses to respond to *P. aeruginosa* antagonism: inactivation of the *P. aeruginosa* siderophore pyochelin via methylation. Using a combination of mass spectrometry imaging (MSI), classical molecular networking, mass spectrometry guided screening, genetic mutation, and *in vitro* and *in vivo* assays, we show that some *S. aureus* strains encode an uncharacterized methyltransferase, Spm (<u>s</u>taphylococcal <u>p</u>yochelin <u>m</u>ethyltransferase), that neutralizes the siderophore and ROS generating functions of pyochelin. Importantly, presence of Spm is critical for *S. aureus* co-existence with *P. aeruginosa* in a murine wound co-infection model.

## Results

### MRSA methylates pyochelin produced by *P. aeruginosa*

To study the natural product mediated interaction of *S. aureus* and *P. aeruginosa,* USA300 methicillin-resistant *S. aureus* strain LAC (MRSA) and *P. aeruginosa* strain PAO1 (PAO1) were grown as mono-cultures or adjacent co-cultures on tryptic soy agar (TSA). After 24 h growth at 37°C, significant inhibition of the MRSA colony by PAO1 was observed when the two bacterial species were co-cultured (Figures 1A and S1A). A section of the agar was excised from the mono- and co-cultures and examined for the spatial distribution of the secreted metabolites by matrix-assisted laser desorption ionization mass spectrometry imaging (MALDI-MSI)^40,41^. Although dozens of ions were observed, the ion *m/z* 339, was exclusively detected in the co-cultures and predominantly distributed towards the MRSA colony, albeit at low intensity (Figure 1A). As this ion was not detected in PAO1 or MRSA mono-cultures, we hypothesized that it was produced by MRSA and the observed low signal intensity was likely due to the low MRSA cell mass in the interaction due to inhibition by PAO1.

**Figure 1:**
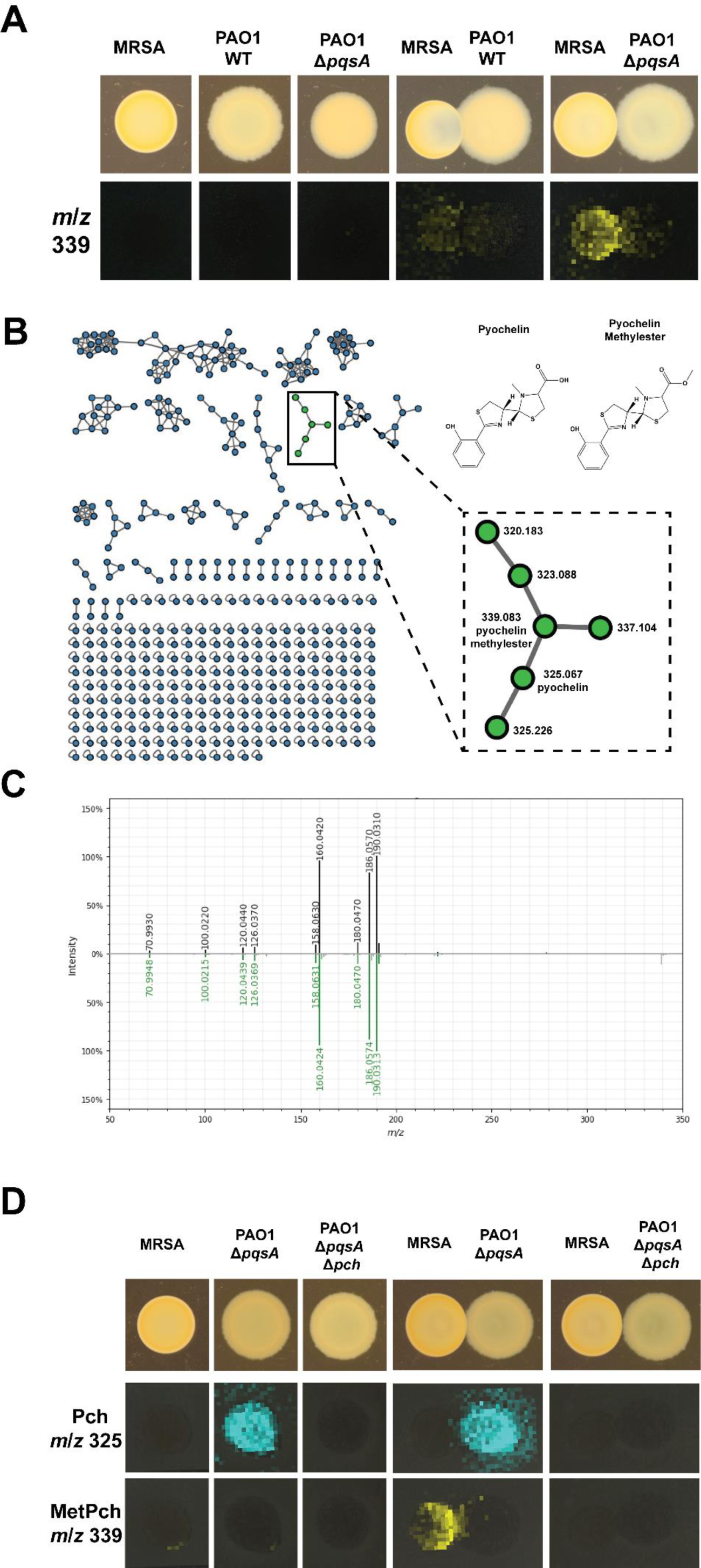
Pyochelin methylester (MetPch) is produced during interaction of *S. aureus* and *P. aeruginosa* **A)** MALDI-MSI of *S. aureus* and *P. aeruginosa* grown as mono cultures or interactions. Photographs of the cultures are shown on top. *P. aeruginosa* wildtype (PAO1 WT) but not a *P. aeruginosa* quinolone mutant (PAO1 Δ*pqsA*) inhibits *S. aureus* wildtype (MRSA) when grown in co-culture. The bottom row shows the false colored *m/z* distribution of an unknown compound with *m/z* 339 that was exclusively observed during interaction of *P. aeruginosa* and *S. aureus*. **B**) The interaction specific compound (*m/z* 339) is annotated as pyochelin methylester (MetPch), a member of the pyochelin (Pch) molecular family, by molecular networking. **C)** Mirror plot comparing the MS^2^ spectrum of isolated MetPch (top; black trace) with its GNPS library match (bottom; green trace). **D)** MALDI-MSI of *S. aureus* and *P. aeruginosa* producing (Δ*pqsA*) or not producing pyochelin (Δ*pqsA* Δ*pch*). Photographs of the cultures are shown on top. The middle row shows the false colored *m/z* distribution for Pch (*m/z* 325) and the bottom row for MetPch (*m/z* 339). Pch production can be observed by PAO1 Δ*pqsA*, but not PAO1 Δ*pqsA* Δ*pch*.

*P. aeruginosa* produces a variety of secreted metabolites that influence *S. aureus* growth directly and indirectly, including the alkyl quinolones. 2-heptyl-4(1H)-quinolone (HHQ) and 2-heptyl-3-hydroxy-4(1H)-quinolone (*<u>P</u>seudomonas* <u>q</u>uinolone <u>s</u>ignal; PQS) regulate the production of anti-staphylococcal factors, such as pyocyanin and rhamnolipids, while 2-heptyl-4-quinolone-N-oxide (HQNO) directly suppresses *S. aureus* growth by inhibiting the respiratory chain^36,45–47^. The *pqsABCDE* biosynthetic gene cluster is required for *P. aeruginosa* to produce these metabolites. Therefore, the gene encoding PqsA, an anthranilate coenzyme A ligase that catalyzes the first step in alkyl quinolone biosynthesis, was deleted to render PAO1 (PAO1 Δ*pqsA*) deficient in the production of alkyl quinolones and pyocyanin and impair MRSA inhibition^42–45^. A replicate interaction of MRSA with PAO1 Δ*pqsA* was inoculated and subjected to MALDI-MSI. As expected, abrogation of quinolone production by the PAO1 Δ*pqsA* strain (Figures 1A **and S1A, top panels**) led to increased MRSA growth with a concomitant increase in signal intensity for the ion at *m/z* 339 compared to the wild-type interaction (Figure 1A).

To facilitate the identification of the ion at *m/z* 339, untargeted liquid chromatography high resolution tandem mass spectrometry (LC-MS/MS) metabolomics data were collected on a chemical extraction of a duplicate interaction of MRSA and PAO1 Δ*pqsA* and the data were analyzed using the classical molecular networking workflow of the GNPS platform^46^. Classical molecular networking organizes compounds into molecular families by using MS/MS spectral relatedness as a proxy for structural similarity. Within the molecular network, MS/MS spectra are represented as nodes connected by edges that indicate spectral similarity. The MS/MS spectra in the data analyzed are simultaneously scored for spectral similarity to the data within the GNPS spectral libraries to provide putative annotation of compounds.

In the molecular network generated, the node representing *m/z* 339.0829 clustered within the pyochelin molecular family^35^ (Pch, [M+H]+, calc 325.0681, *m/z* 325.0671, −2.95 ppm) and was putatively annotated as pyochelin methyl ester, where methylation of the Pch structure was localized to the carboxylic acid group (MetPch, [M+H]+, calc: 339.0837, m/z 339.0829, −2.43 ppm). (Figures 1B and 1C). This structural annotation of pyochelin and pyochelin methyl ester was confirmed by comparing the retention time, exact mass, and MS/MS fragmentation of the microbially produced metabolites to synthesized standards (**Figures S1B and S1C**).

Based upon the spatial distribution of MetPch in the MSI data, we hypothesized that *S. aureus* methylates Pch produced by *P. aeruginosa* by an unknown mechanism. This hypothesis is supported by the fact that *P. aeruginosa* is a known producer of Pch^47^, while *S. aureus* does not harbor biosynthetic genes for the production of Pch or related molecules. To verify that production of MetPch was dependent upon Pch biosynthesis by *P. aeruginosa*, MRSA was co-cultured with the PAO1 *pqsA* mutant impaired in Pch biosynthesis (PAO1 Δ*pqsA* Δ*pch*) on TSA. Indeed, absence of MetPch in the co-culture was associated with the loss of Pch production by PAO1 Δ*pqsA* Δ*pch* (Figure 1D).

Although the spatial distribution of MetPch favored localization to the MRSA colony, several CF lung *P. aeruginosa* isolates produce a methylated pyochelin^48,49^. To confirm that MRSA was the biosynthetic origin of MetPch during interaction with PAO1, MRSA cultures were incubated with cell free supernatant from PAO1 Δ*pqsA*, which contains Pch (**Figure S2A**). While low levels of MetPch were detected in PAO1 Δ*pqsA* cell free supernatant, incubation of the supernatant with MRSA increased the levels of MetPch over 170-fold (**Figure S2B**). It is important to note that the steep increase of MetPch levels in PAO1 Δ*pqsA* supernatant treated with MRSA compared to the untreated PAO1 Δ*pqsA* supernatant control did not correspond to an equivalent decrease in Pch in the MRSA treated PAO1 Δ*pqsA* supernatant (**Figure S2A**). Quantification of synthetic Pch and MetPch under the same conditions resulted in an approximately 10-fold higher signal intensity of MetPch compared to Pch at equimolar concentrations (**Figure S2C**) suggesting a difference in the ionization efficiency of the two compounds during electrospray ionization (ESI). Taken together, these results indicate that MRSA likely produces a methyltransferase that methylates the carboxylic acid of Pch to produce MetPch.

### Pyochelin is methylated by <u>S</u>taphylococcal <u>p</u>yochelin <u>m</u>ethyltransferase (Spm)

To identify the *S. aureus* gene encoding the enzyme responsible for biotransformation of Pch to MetPch, an LC-MS guided screen of the Nebraska Transposon Mutant Library^50^, a fully sequenced and sorted library of transposon insertions in the USA300 *S. aureus* LAC derivative JE2, was employed to identify mutants that did not produce MetPch when incubated with cell free PAO1 Δ*pqsA* supernatant containing Pch. Of 576 mutants tested, transposon mutant NE502 (transposon inserted in gene SAUSA300_1977) was the only one unable to produce MetPch (**Supplementary Data 1 - 6**). *In silico* analysis of the SAUSA300_1977 gene sequence identified it as encoding a putative methyltransferase (TcmP-like; IPR016874), which we termed <u>S</u>taphylococcal <u>p</u>yochelin <u>m</u>ethyltransferase (Spm).

To assess whether Spm homologues were present within other staphylococcal genomes, the sequence of the *spm* gene was searched against the genomes available in the KEGG database (Figure 2A). This analysis revealed that Spm is conserved in 17 out of 53 *S. aureus* strains (32%) and in 4 out of 35 coagulase negative staphylococci (11%) represented in the database. To substantiate that strains that encode *spm* in their genomes were capable of converting Pch to MetPch, six *S. aureus* strains with sequenced genomes (including LAC USA300) were incubated with Pch and evaluated for MetPch production using LC-MS. The four *S. aureus* strains harboring an *spm* homologue, including USA300 LAC, Newman, COL and MW2, produced MetPch, while the two *S. aureus* strains lacking *spm* (UAMS-1 and MN8) did not (Figures 2B and 2C).

**Figure 2:**
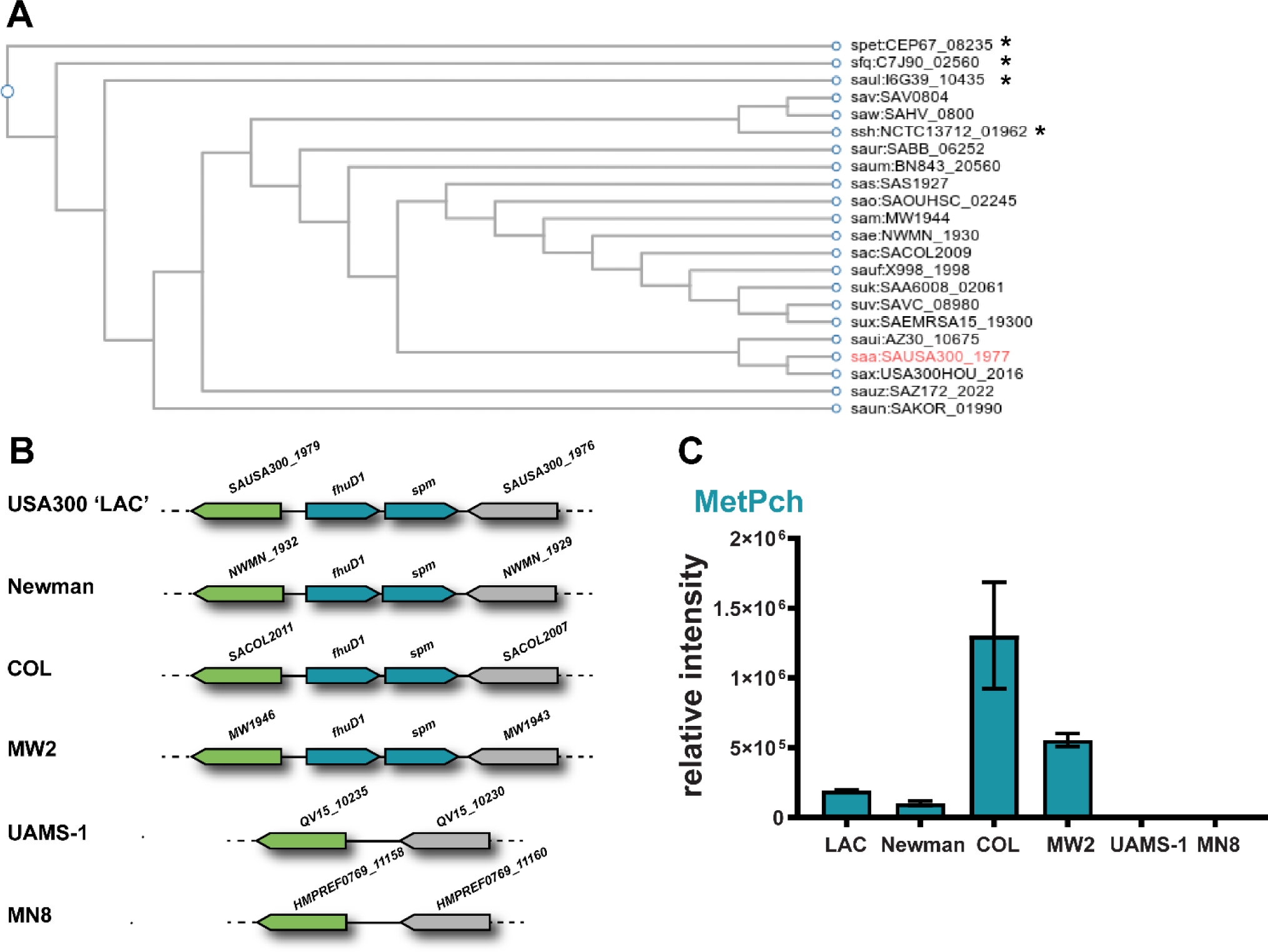
Pyochelin methyl ester is only produced by *S. aureus* strains harboring *spm*. **A)** Dendrogram of Spm (SAUSA300_1977) orthologs in staphylococcal species. Dendrogram was constructed based on KEGG database entries. *coagulase-negative staphylococcal species. **B**) Genetic environment of *spm* (SAUSA300_1977) in six sequenced *S. aureus* strains. The gene *spm* is encoded in a putative operon with *fhuD1*. **C)** MetPch was detected from cultures of *S. aureus* strains encoding the *spm* gene but not by *spm* negative strains (UAMS-1, MN8) when incubated with PAO1 Δ*pqsA* supernatant . n = 4 biological replicates. Values are mean ± SDs.

To further establish that Spm is responsible for the methylation of Pch, a markerless in-frame deletion of *spm* in the MRSA background (MRSA Δ*spm*) was generated and this mutant was incubated with cell free supernatant from PAO1 Δ*pqsA*, which contains Pch. Compared to MRSA wild type (WT), MRSA Δ*spm* produced significantly lower levels of MetPch, while genetic complementation of MRSA Δ*spm* with *spm* (MRSA Δ*spm*::*spm*) recovered production of MetPch to WT levels (Figure 3A). These results confirm that Spm is necessary for the efficient biotransformation of Pch to MetPch by *S. aureus*.

**Figure 3:**
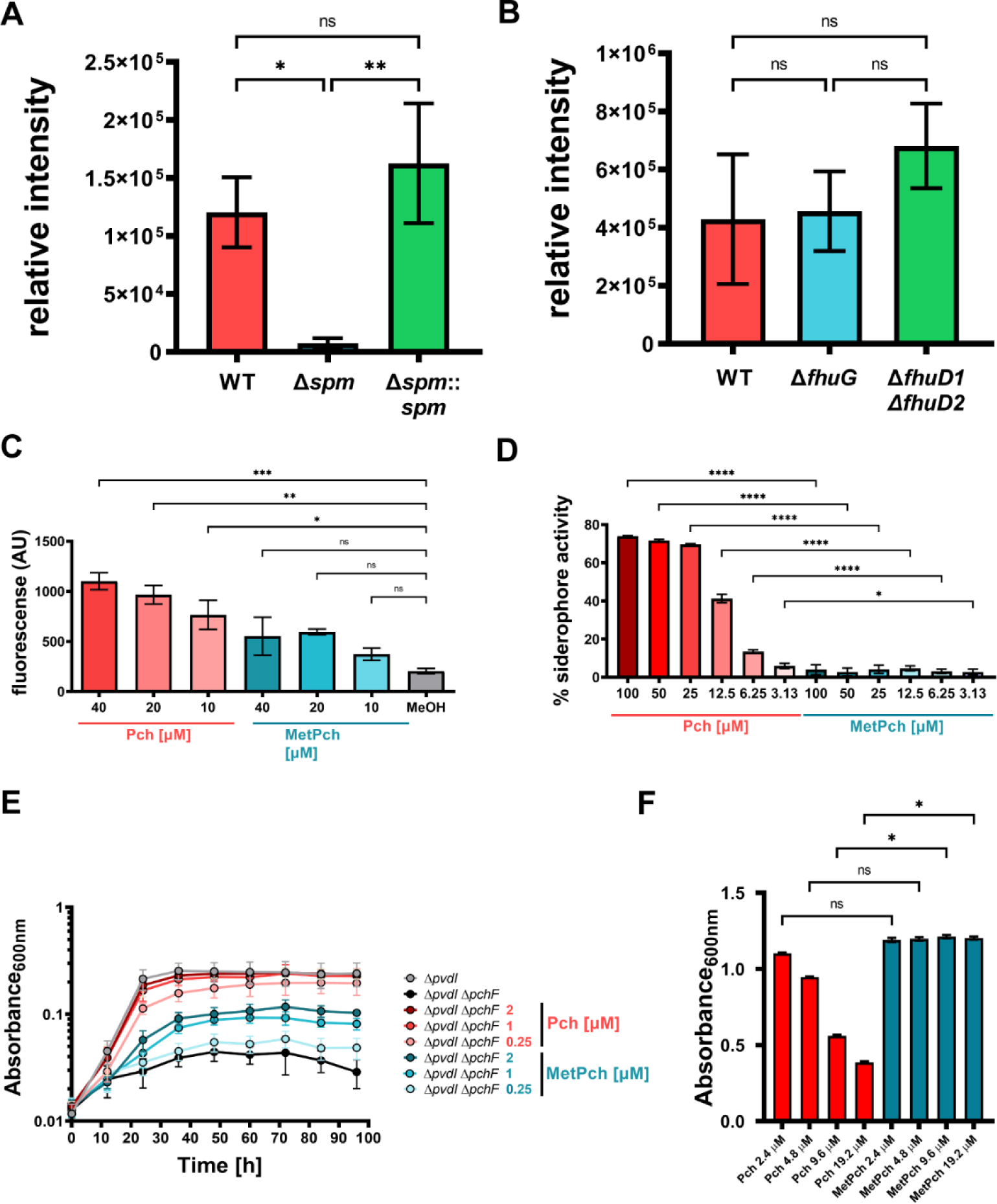
Spm mediated methylation leads to Pch inactivation. **A)** Incubation of PAO1 Δ*pqsA* cell free supernatant with overnight cultures of MRSA wildtype (WT), an *spm* mutant strain (Δ*spm*) and its corresponding complementation strain (Δ*spm*::*spm*) results in high amounts of MetPch production by WT and Δ*spm*::*spm*, but not by Δ*spm*. n = 3 biological replicates. [one-way ANOVA with Tukey’s multiple comparison] **B)** The Fhu xenosiderophore uptake machinery does not promote MetPch production. Mutation of the two xenosiderophore receptors *fhuD1* and *fhuD2 (*Δ*fhuD1* Δ*fhuD2)* but not *fhuG* (Δ*fhuG*), which is essential for the uptake of several hydroxamate xenosiderophores, does result in significantly lowered MetPch production. n = 3 biological replicates. [one-way ANOVA with Tukey’s multiple comparison] **C)** Exposure to Pch but not MetPch results in significantly elevated ROS production of WT compared to the carrier control (MeOH). n = 3 biological replicates. [Kruskal-Wallis with Dunn’s multiple comparison] **D)** Siderophore activity of Pch and MetPch were evaluated by liquid Chrome Azural S assay. Siderophore activity of MetPch is significantly reduced compared to Pch across all tested concentrations. n = 5 biological replicates. [one-way ANOVA and Šidák’s multiple comparison] **E)** Pch but not MetPch supports *P. aeruginosa* growth under low iron conditions. 96-hour growth curves of a PAO1 pyoverdine mutant (Δ*pvdI*) and a pyoverdine/pyochelin double mutant (Δ*pvdI* Δ*pchF*) supplemented with different concentrations of synthetic Pch or MetPch. Cultures were grown in M9 minimal medium with glycerol as carbon source. To create a low iron environment, 500 µM 2,2’- Bipyridine was added to the growth medium. n = 5 biological replicates from 4 independent experiments. **F**) Exposure to Pch but not MetPch results in growth inhibition of MRSA under low iron conditions (chelex treated TSB). Growth was significantly reduced in samples treated with high concentrations of Pch (9.6 µM and 19.2 µM) compared to samples treated with equimolar amounts of MetPch. n = 3 biological replicates. [Kruskal-Wallis with Dunn’s multiple comparison] **B – D, E)** Values are mean ± SDs. \**P* ≤ 0.05, \*\**P* ≤ 0.01, \*\*\**P* ≤ 0.001, \*\*\*\**P* ≤ 0.0001, ns = not significant.

Spm is encoded in a putative operon downstream of the xenosiderophore receptor FhuD1 ^51^. The genomes of *S. aureus* strains that do not encode the *spm* gene lack the whole *fhuD1*- *spm* locus (Figure 2B). FhuD1 and FhuD2 are xenosiderophore receptors that bind iron (III)- hydroxamate siderophore complexes, with a partially overlapping substrate range, that function as part of the ferrichrome uptake (Fhu) system^51^. The xenosiderophores bound by these two receptors are transported into *S. aureus* cells via the siderophore-permeases and ATPase transporter machinery encoded by the *fhuCBG* operon^52^. Although Pch is a phenolate siderophore and the Fhu xenosiderophore acquisition system is used by *S. aureus* to acquire iron from exogenous hydroxamate siderophores, the proximity of *spm* to *fhuD1* within the MRSA genome suggested that the Fhu system may directly or indirectly influence methylation of Pch. To test the influence of the Fhu system on the production of MetPch, markerless in-frame deletions of *fhuD1*/*fhuD2* and *fhuG* in the MRSA background were generated (MRSA Δ*fhuD1* Δ*fhuD2* and MRSA Δ*fhuG,* respectively) and these mutants were incubated with PAO1 Δ*pqsA* cell free supernatant containing Pch. Although MRSA Δ*fhuD1* Δ*fhuD2* produced higher levels of MetPch than MRSA WT and MRSA Δ*fhuG,* the difference in production levels between the strains was not statistically significant (Figure 3B).

The ferric uptake regulator (Fur) is an iron-dependent transcriptional repressor that regulates genes involved in iron import and acquisition^53^. When iron is abundant, Fur binds ferrous iron and blocks transcription of iron acquisition genes. The promoter region of the *fhuD1*/*spm* operon contains a fur box^51^, a conserved sequence for Fur-binding, indicating that this operon may be regulated by Fur activity. To measure transcriptional activation of the *fhuD1*/*spm* operon, sGFP was fused to the *fhuD1*/*spm* promoter and fluorescence was measured in both wild type MRSA and *fur* mutant (MRSA *fur*::*tet*). Promoter activity was significantly increased in MRSA *fur*::*tet* compared to WT (**Figure S2D**), indicating that Fur represses its activity and production of MetPch is likely Fur-dependent. Collectively, these results suggest that *S. aureus* produces Spm to methylate Pch in low iron environments.

### Methylation of Pch neutralizes its biological functions

Pch is a phenolate siderophore produced by *Pseudomonas* and *Burkholderia* species^35,54^. In addition to its iron chelating activity^47^ , exposure to Pch induces intracellular reactive oxygen species (ROS) production in several bacterial species, including *S. aureus*^36,37^. We hypothesized that methylation of the carboxylic acid of Pch by *S. aureus* could disrupt Pch- driven biological activities.

To test if MetPch had reduced antibacterial activity against *S. aureus* compared to Pch, MRSA was treated with equimolar concentrations of synthetic Pch and MetPch (10, 20 and 40 µM) and ROS levels were assessed by measuring the production of 2′,7′- dichlorofluorescein (DCF) from 2′,7′-dichlorodihydrofluorescein (H2DCF) by fluorescence (Figure 3C). While MRSA treated with Pch had a concentration-dependent increase in ROS production, exposure to MetPch did not result in significant increase in ROS production for all tested concentrations compared to carrier control (Figure 3C).

*P. aeruginosa* produces two siderophores: pyoverdine (Pvd) and Pch. Pvd is considered the primary siderophore of *P. aeruginosa* due to its higher affinity for iron^55^. However, Pch is produced first and Pvd is only produced at extremely low concentrations of iron, likely due to the metabolic burden of its production^34^. Importantly, the carboxylic acid moiety of Pch is required for iron binding as well as its interactions with its receptor (FptA) located on the outer membrane of *P. aeruginosa*^56^. To determine if MetPch had reduced iron binding capacity compared to Pch, iron binding of synthetic Pch and MetPch was measured via Chrom Azurol S (CAS) assay. Iron binding by MetPch was significantly lower than iron binding by Pch for all concentrations tested, ranging from 3.13 µM to 100 µM (Figure 3D).

To ascertain if the reduced iron binding of MetPch observed in the CAS assay reflected reduced support for *P. aeruginosa* growth under iron-limited conditions, a PAO1 mutant strain deficient in the production of both Pvd and Pch (PAO1 Δ*pchF* Δ*pvdI,*) was generated and chemically complemented with synthetic Pch or MetPch at equimolar concentrations in low iron medium (M9 minimal medium supplemented with 500 µM 2,2′-Bipyridine) (Figure 3E). Only PAO1 Δ*pchF* Δ*pvdI* is unable to grow robustly under iron-limited conditions as the production of either siderophore is sufficient to support *P. aeruginosa* growth (**Figure S3**). While chemical complementation of PAO1 Δ*pchF* Δ*pvdI* with Pch was sufficient to recover *P. aeruginosa* growth to PAO1 Δ*pvdI* levels, MetPch was not (Figure 3E). Conversely, when *S. aureus* was grown in low iron medium (chelex treated TSB) and subjected to Pch or MetPch, high concentrations of Pch inhibited its growth, while MetPch had no effect on *S. aureus* growth at any of the concentrations tested (Figure 3F). These results indicate that methylation of the carboxylic acid of Pch by *S. aureus* abrogates iron binding and ROS generation by Pch, providing a protective effect from this compound.

### Spm increases *S. aureus* fitness in co-infection with PAO1

Taken together, our *in vitro* results show that *S. aureus* produces Spm to methylate the carboxylic acid moiety of Pch to protect itself from ROS production and iron limitation during interaction with *P. aeruginosa*. This ability of *S. aureus* to diminish Pch bioactivity may represent a viable competition strategy for *S. aureus* to co-exist with *P. aeruginosa* in skin wounds, one of their primary interaction environments. To test this hypothesis, a murine skin wound model was used to evaluate PAO1 and MRSA virulence and fitness as well as microbial competition between PAO1 and MRSA, MRSA Δ*spm*, or MRSA Δ*spm*::*spm* (Figure 4A).

**Figure 4:**
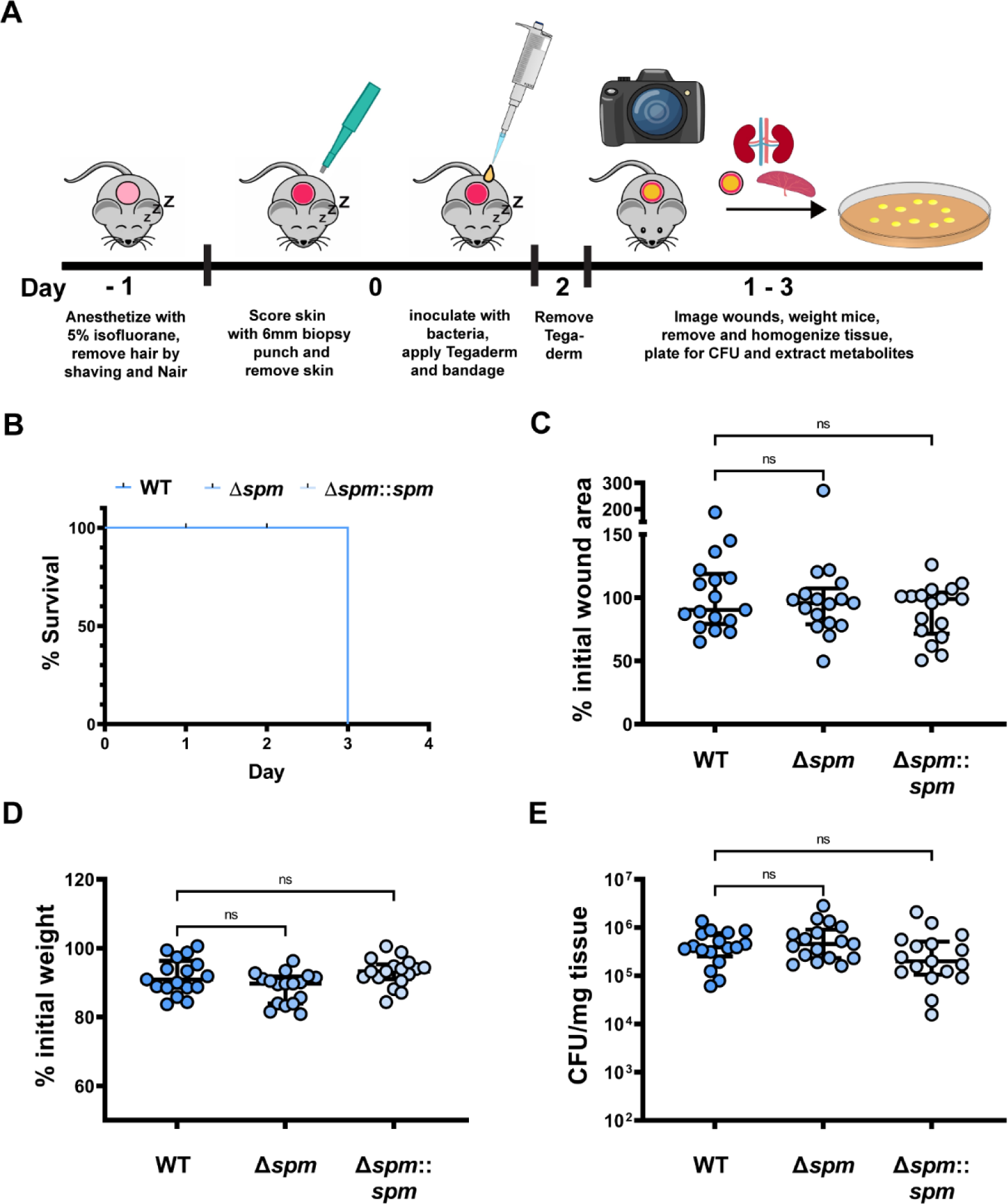
Spm does not change fitness or virulence of MRSA. **A**) Schematic of our murine wound skin infection model. **B**). Murine survival after skin wound infection with MRSA wild type (WT), the *spm* mutant strain (Δ*spm*) and the *spm* complemented mutant (Δ*spm*::*spm*). **C**) The change in murine skin wound size area between start and end of the experiment. Changes were calculated from the measured wound size area before bacterial infection and at the endpoint of the murine skin wound model. The changes are expressed as percent (%) of the initial wound size area at the endpoint of the experiment. **D**) The change in murine weight between start and end of the experiment. Changes were calculated from the murine weight before bacterial infection and at the endpoint of the murine skin wound model. The changes are expressed as percent (%) of the initial murine weight at the endpoint of the experiment. **E**) MRSA colony forming units (CFU) recovered from mouse tissue at the endpoint of the experiment. **B – E**) n = 17 biological replicates from 3 independent experiments. **C – E**) [Kruskal-Wallis with Dunn’s multiple comparison]. Values are median ± interquartile range. Each dot represents values from a single mouse. ns = not significant.

Spm is an uncharacterized enzyme and we first measured its influence on MRSA wound infections. Single infections with MRSA WT, MRSA Δ*spm*, and MRSA Δ*spm::spm* were performed to determine if Spm influences MRSA fitness or infection progression. Murine survival as well as changes to wound size and murine weight between the day of infection and the day of sacrifice were assessed as indicators of infection severity. All mice from the three groups (MRSA WT, MRSA Δ*spm*, and MRSA Δ*spm::spm)* survived over the course of the three day experiment and no significant changes in wound size, weight or MRSA fitness, as determined by colony forming unit (CFU) recovery of the three MRSA strains, were observed (Figures 4B to 4E). This demonstrates that Spm does not affect the ability of MRSA to cause infection or to persist in our *in vivo* murine skin wound model.

Pyochelin biosynthesis by PAO1 in the wound environment is a prerequisite to assess the role of Spm during bacterial wound co-infection. Therefore, we established a protocol for pyochelin extraction from PAO1 wound infections and successfully quantified Pch produced by PAO1 in our murine skin wound infection model (Figure 5A). No quantifiable MetPch amounts were detected in PAO1 mono infections, but examination of our raw data showed that several samples contained trace amounts of MetPch that fell below the limit of quantitation threshold for our data analysis workflow.

**Figure 5:**
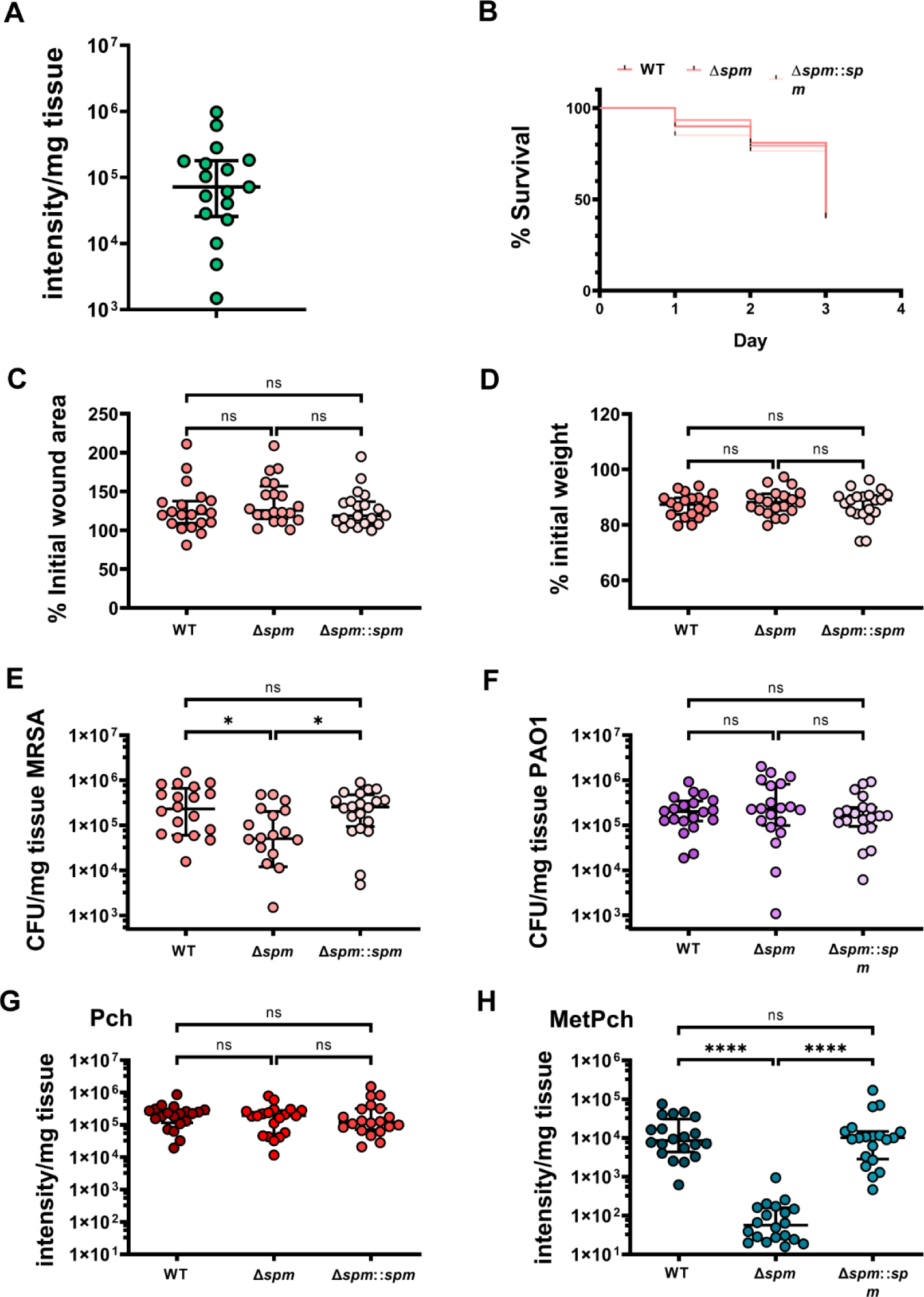
Mutation of *spm* impairs the fitness of MRSA during co-infection with PAO1. **A**) Quantification of Pch recovered from mouse tissue infected with PAO1. **B**) Murine survival after skin wound co-infection with PAO1 and MRSA. **C**) The change in murine skin wound size area between start and end of the experiment. Changes were calculated from the measured wound size area before bacterial infection and at the endpoint of the murine skin wound model. The changes are expressed as percent (%) of the initial wound size area at the endpoint of the experiment. **D**) The change in murine weight between start and end of the experiment. Changes were calculated from the murine weight before bacterial infection and at the endpoint of the murine skin wound model. The changes are expressed as percent (%) of the initial murine weight at the endpoint of the experiment. **E**) MRSA and **F**) PAO1 colony forming units (CFU) recovered from mouse tissue at the endpoint of the experiment. Quantification of **G**) Pch and **H)** MetPch recovered from mouse tissue co-infected with PAO1 and MRSA. All co-infections were performed with PAO1 wild type and either MRSA wild type (WT), the Spm mutant strain (Δ*spm*) or the Spm complemented strain (Δ*spm*::*spm*). The co- infecting MRSA strain is indicated in the figure legend **B**) or below the X-axis **C - H**). **A**) n = 17 biological replicates from 3 independent experiments. **C – H**) n = 20 biological replicates from 4 independent experiments. [Kruskal-Wallis with Dunn’s multiple comparison]. Values are median ± interquartile range. Each dot represents values from a single mouse. \**P* ≤ 0.05, \*\**P* ≤ 0.01, \*\*\**P* ≤ 0.001, \*\*\*\**P* ≤ 0.0001, ns = not significant.

Next, our murine wound co-infection model was employed to determine the role of Spm during MRSA/PAO1 wound co-infection. Infection severity was assessed by the same indicators previously used for the MRSA single infections (murine survival, wound size progression, weight loss). Mice co-infected with MRSA WT and PAO1 WT showed a lowered, although not significant, probability to survive until the end of the experiment (day 3) compared to mono infections with MRSA WT or PAO1 WT (**Figure S4A**). In addition, a significant increase in wound size progression in mice co-infected with MRSA WT and PAO1 WT was observed compared to mono infections (**Figure S4B**) and significantly less weight loss in mice infected with MRSA WT compared to PAO1 WT but not compared to co- infections (**Figure S4C**).

Comparison of the infection severity outcomes of the three co-infections of PAO1 with either MRSA, MRSA Δ*spm*, or MRSA Δ*spm*::*spm* showed no difference in murine survival, wound size progression, or weight loss (Figure 5B-D). However, MRSA fitness was significantly altered between Spm positive and Spm negative MRSA as evidenced by the significantly reduced recovery of colony forming units (CFUs) of MRSA Δ*spm* compared to MRSA WT and MRSA Δ*spm*::*spm* from the wounds (Figure 5E). *P. aeruginosa* fitness was similar for all co-infections (Figure 5F). Collectively, our data indicates that Spm is not involved in MRSA virulence but is required for optimal fitness of MRSA in polymicrobial skin wound infection with *P. aeruginosa*.

To determine if MRSA produced MetPch from Pch synthesized by PAO1 *in vivo*, Pch and MetPch were quantified from wound tissues of the murine skin wound co-infections. Both Pch and MetPch were detected in the wound tissue. While comparable amounts of Pch were measured in all three co-infections (Figure 5G), MetPch was produced at more than 100-fold lower in MRSA Δ*spm* co-infections with PAO1 compared to co-infections with MRSA WT and these levels were recovered with genetic complementation of the *spm* mutant strain (Δ*spm*::*spm*) (Figure 5H). Importantly, levels of MetPch positively correlated to recovered MRSA CFUs in the polymicrobial skin wound infection model (Spearman’s rank correlation coefficient; ρ = 0.5186), further supporting that *S. aureus* deactivates Pch through Spm- mediated methylation to enhance fitness in the presence of *P. aeruginosa* (Figure 5E), which likely contributes to co-existence in these dual species infections.

## Discussion

Using a mass spectrometry directed approach, we investigated the interaction of *S. aureus* and *P. aeruginosa in vitro* and in an *in vivo* murine skin wound model that reflects the early stages of wound co-infection. We demonstrate an underappreciated role for the *P. aeruginosa* derived siderophore pyochelin during *in vivo* competition of PAO1 and MRSA in skin wound co-infection. In addition, our data clearly shows that *S. aureus* inactivates pyochelin by methylation via the previously uncharacterized enzyme Spm and this translates to enhanced fitness of MRSA in a wound co-infection model with PAO1 (Figure 6). We validated that the methylation of Pch by Spm occurs on the carboxylic acid group by comparing the experimentally-derived compounds with chemically synthesized authentic standards.

**Figure 6:**
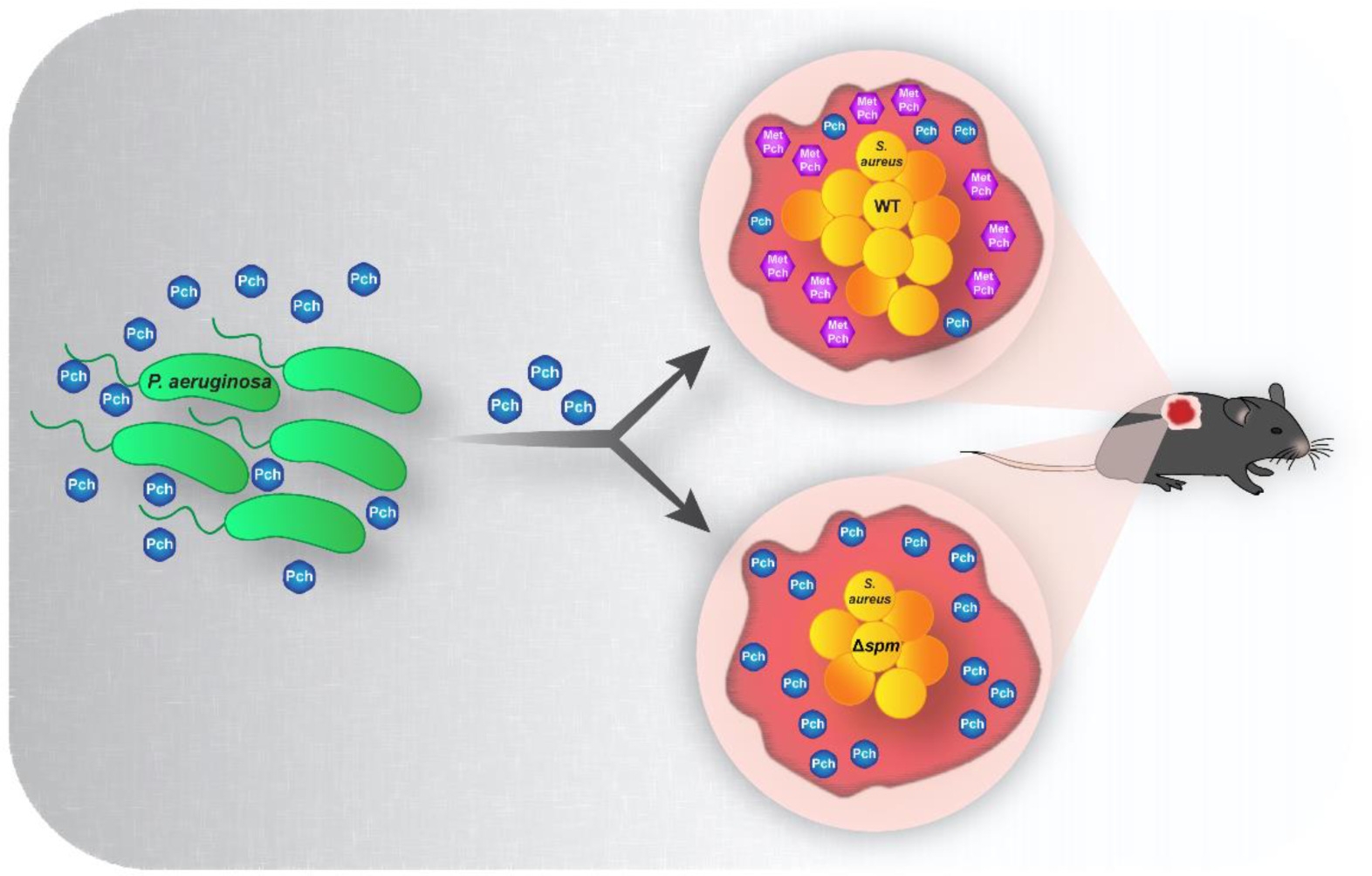
Model of *S. aureus*/*P. aeruginosa* chemical interaction. *P. aeruginosa* produces and secretes pyochelin during infection. *S. aureus* wild type cells inactivate pyochelin through enzymatic methylation. *S. aureus* mutant cells lacking the Spm methyltransferase are unable to methylate pyochelin. Pyochelin methylation leads the loss of siderophore activity and reduced intracellular ROS production in *S. aureus* cells, resulting in a higher fitness of *S. aureus* during competition with *P. aeruginosa in vivo*.we

The *spm* gene is encoded in an operon with the *fhuD1* gene, that encodes a xenosiderophore receptor involved in the uptake of ferrichrome and ferrioxamine B^51^, suggesting a possible role of the Fhu xenosiderophore uptake machinery in pyochelin methylation. Simultaneous mutation of *fhuD1* and *fhuD2* resulted in elevated MetPch production, but mutation of *fhuG*, an essential component of the Fhu xenosiderophore import machinery, did not result in any change in MetPch production. These data indicate that the Fhu system is most likely not involved in Pch or MetPch transportation, but that FhuD1 and FhuD2 might bind Pch and exclude it from Spm-mediated methylation. Alternatively, FhuD1 and FhuD2 might bind MetPch, preventing its diffusion into the cell free supernatant and exclude it from our metabolite extracts.

We show that methylation of Pch by Spm contributes to enhanced MRSA fitness in a skin wound co-infection model with *P. aeruginosa*. Based upon the similarity of the infection outcomes, including survival, wound size progression and weight loss, in the mono-species infection model with WT MRSA and a deletion mutant of Spm, we determined that Spm does not contribute to MRSA virulence in our murine skin wound infection model. However, in the co-infection model with PAO1, Spm was required to maintain WT population levels of MRSA. From the two-dimensional distribution of MetPch in the MALDI-MSI data, we postulate that methylation of Pch is localized to *S. aureus* growth *in vivo*. From our *in vitro* studies, local methylation of Pch by *S. aureus* would result in increased survival by impeding iron binding by Pch, thereby improving its ability to compete for environmental iron, and decreasing induction of its own ROS by detoxifying Pch.

Intriguingly, deactivation of Pch by esterification of the carboxylic acid moiety appears to be a conserved mechanism between a variety of microorganisms, including the fungus *Phellinus noxius*^57^ and soil bacterium *Bacillus amyloliquefaciens*^58^. The apparent wide-spread mechanism by microbes from different kingdoms to deactivate Pch via chemical modification suggests that Pch may have a larger impact on microbial ecology than currently appreciated. The herein presented study is the first to demonstrate the molecular and genetic mechanism behind pyochelin methylation and its importance for *S. aureus* fitness in polymicrobial infection.

## Methods

### Media and growth conditions

*Staphylococcus aureus, Pseudomonas aeruginosa,* and *Escherichia coli* strains and plasmids used in this work are listed in **Table S1**. *S. aureus* was regularly cultured in Bacto Tryptic Soy Broth (TSB; BD) and *E. coli* and *P. aeruginosa* were cultured in Lennox Lysogeny Broth (LB; RPI) at 37 °C with 200 rpm shaking. To create environments with low iron availability, media were treated with Chelex-100 (chelex: BioRad) as follows: 0.5 % (w/v) Bacto Casamino Acids Technical (CAA; BD) was incubated for 5 h with 5 g/L chelex, filtered with a Nalgene Rapid-flow Filter Unit (0.2 µm aPES membrane; Thermo Scientific) to remove the chelex resin, adjusted to pH 7.4, autoclaved, and supplemented with MgCl_2_ (0.4 mM final concentration). TSB was incubated with 2 g/L chelex for 3 h, filtered with a Nalgene Rapid-flow Filter Unit (0.2 µm aPES membrane; Thermo Scientific) to remove the chelex resin, and autoclaved. Experiments to determine *P. aeruginosa* growth support by pyochelin and pyochelin methyl ester were performed in M9 minimal medium supplemented with 500 µM 2,2’-Bipyridine (Fisher Scientific). M9 minimal medium was prepared as follows: 5x M9 salts: 64 g/L Na_2_HPO_4_*7H_2_O, 15 g/L KH_2_PO_4_, 2.5 g/L NaCl, and 5g/l NH_4_Cl. 5x M9 salts were diluted in H_2_O containing MgSO_4_ (2 mM final concentration), CaCl_2_ (0.1 mM final concentration), and glycerol (0.5 % (v/v) final concentration). Antibiotics were added to the media where necessary at the following concentrations: chloramphenicol (Cam), 10 μg/mL; erythromycin (Erm), 5 μg/mL; and tetracycline (Tet), 1 μg/mL. *E. coli* strains with plasmids were maintained on media supplemented with ampicillin (Amp) at 100 μg/mL; kanamycin (Kan), 50 μg/mL; gentamicin (Gen) 20 μg/mL or spectinomycin (Spec) at 50 μg/mL.

### Construction of gene deletion mutants

Markerless gene mutations were introduced using the temperature-sensitive plasmid pJB38 for *S. aureus* or pEX2G for *P. aeruginosa* carrying DNA fragments (∼1000 bp in size) flanking the region targeted for deletion. The flanking regions were amplified by Phusion High-Fidelity DNA polymerase (New England BioLabs) with the primers listed in **Table S2**.

#### S. aureus

DNA fragments (∼1000 bp in size) flanking the region targeted for deletion and pJB38 were digested with restriction enzymes, as indicated in **Table S2**, and subsequently purified with the QIAquick PCR purification kit (Qiagen). After triple ligation of the flanking region pairs with pJB38, the resulting plasmid was electroporated into *E. coli* DC10b. *E. coli* cells carrying the plasmid were selected on LB plates containing 100 μg/ml Amp, single colonies were picked, patched on new LB plates containing 100 μg/ml Amp, and the presence of the plasmid containing the flanking regions was confirmed by PCR using the primers listed in **Table S2**. The plasmid was recovered from overnight cultures of positive clones with a QIAquick Spin miniprep kit (Qiagen) and the sequence of the flanking regions was confirmed by in-house sequencing with their respective construction primers and sequencing primers listed in **Table S2**. The plasmid was electroporated into the *S. aureus* target strain, positive clones carrying pJB38 with the desired flanking regions were selected on TSA plates containing 10 μg/ml Cam at 30°C. For homologous recombination, positive clones were streaked on TSA-Cam and incubated at 42°C for 24 h. Cam-resistant colonies were restreaked on TSA-Cam and incubated at 42°C overnight. Single colonies were picked and incubated in 5 ml TSB at 30°C with 200 rpm shaking overnight. The resulting overnight cultures were diluted 1:1000 in TSB and incubated at 30°C with 200 rpm shaking overnight. This was repeated for six consecutive days, and subsequently, dilutions (10^−6^, 10^−7^, and 10^−8^) were plated on TSA plates containing anhydrotetracycline (ahTet) 200 ng/ml for counterselection and incubated at 30°C overnight. Single colonies were then patched on TSA and TSA-Cam plates and grown at 30°C overnight. Plasmid loss was indicated by growth on TSA but not TSA-Cam. Colonies were screened for the desired mutation by PCR with primers (**Table S2**) located on the *S. aureus* chromosome outside of the homology arm regions.

#### P. aeruginosa

DNA fragments (∼1000 bp in size) flanking the region targeted for deletion and pEX2G were digested with restriction enzymes, as indicated in **Table S2**, and subsequently purified with the QIAquick PCR purification kit (Qiagen). After triple ligation of the flanking region pairs with pEX2G, the resulting plasmid was electroporated into *E. coli* Top10. *E. coli* cells carrying the plasmid were selected on LB plates containing 20 μg/ml Gen, single colonies were picked, patched on a new LB plate containing 20 μg/ml Gen, and the presence of the plasmid containing the flanking regions was confirmed by PCR using the primers listed in **Table S2**. The plasmids were recovered from overnight cultures of positive clones with a QIAquick Spin miniprep kit (Qiagen), and the sequence of the flanking regions was confirmed by in-house sequencing with their respective construction primers and sequencing primers listed in **Table S2**. The plasmids were transferred to the *Pseudomonas aeruginosa* PAO1 target strain by conjugation. Single recombinants were selected on Vogel- Bonner minimal medium (VBMM) ^59^ supplemented with 160 μg/ml Gen to create merodiploid strains. The merodiploid strains were resolved by an overnight outgrowth in LB followed by plating on VBMM supplemented with 7.5% sucrose for counterselection. Single colonies were picked, patched on VBMM agar plates containing 7.5% sucrose and gene deletions were confirmed by PCR with primers located on the chromosome outside of the homology arm regions as listed in **Table S2**.

### Construction of complemented *S. aureus* mutant strains

The *spm* mutation in the MRSA Δ*spm* background was repaired by introducing the intact *spm* gene at its original location on the chromosome. The complemented *spm* strain (MRSA Δ*spm*::*spm*) was constructed by amplifying the *spm* gene and flanking regions from the MRSA wild type genome. First, the *spm* gene and a region covering 508 bp upstream of *spm* were amplified with primers listed in **Table S2**, digested with KpnI and SalI and ligated into pJB38 digested with the same restriction enzymes. The resulting plasmid was electroporated into *E. coli* DC10B. After propagation in the *E. coli* strain, the plasmid was digested with NcoI and SalI and ligated with a DNA fragment spanning 559 bp downstream of the *spm* coding sequence which was amplified with primers listed in **Table S2** digested with NcoI and SalI. The resulting plasmid (pJB38*spm*KI) carries the wild type *spm* gene and flanking regions for homologues recombination and an NcoI restriction site located 110 bp downstream of the *spm* stop codon that serves as a watermark for the repaired strain. Introduction of the plasmid into *E. coli* DC10B, transfer to MRSA Δ*spm*, selection for repaired strains and plasmid sequencing was carried out as described for gene deletion mutant construction

### Construction of *S. aureus* LAC *fur* mutant

The *fur*::tet cassette from AH2367 was transduced into *S. aureus* USA300 LAC using phage, and deletions were confirmed by PCR with primers HC479 and HC480 (**Table S2**)

### Construction of promoter fusion vector

A transcriptional fusion of the *fhuD1*/*spm* promoter to sGFP was constructed using plasmid pHC48. A region containing the putative promoter region of the *fhuD1*/*spm* operon was amplified by Phusion High-Fidelity DNA polymerase (New England BioLabs) with the primers listed in **Table S2**. The PCR products and pHC48 were digested with restriction enzymes, as indicated in **Table S2**, and subsequently purified with the QIAquick PCR purification kit (Qiagen). The resulting plasmid, pHC48-spmprom, was electroporated into *E. coli* DC10b. *E. coli* cells carrying the plasmid were selected on LB plates containing 10 μg/ml Cam, single colonies were picked, patched on a new LB plate containing 10 μg/ml Cam, and the presence of the plasmid containing the promoter region was confirmed by PCR using the primers listed in **Table S2**. The plasmid was recovered from overnight cultures of positive clones with a QIAquick Spin miniprep kit (Qiagen), and the sequence of the flanking regions was confirmed by in-house sequencing with their respective construction primers and sequencing primers listed in **Table S2**.

### Promoter fusion activity

*S. aureus* cells harboring the *spm* promoter fusion vector pHC48-spmprom were grown overnight in TSB containing Cam 10 µg/ml or Cam 10 µg/ml and Tet 2 µg/ml for the *S. aureus* LAC *fur*::tet background. Cells were washed with PBS and resuspended in fresh media. Cultures were set to an OD_600_ of 0.05 and 200 µl of each sample were transferred to 96-well black plates with clear bottom (Corning Incorporated) and incubated at 37 °C with 1,000 rpm (Microtiter Plate Shaker Incubator SI505; Stuart) for 24 h. Promoter activities were measured with an Infinite M PLEX plate reader (Tecan) with the following settings: excitation wavelength 549 nm, emission wavelength 588 nm, number of flashes 10, settle time 10 ms, gain 80, integration time 40 µs, Z-position 20,000 µm. To assure similar growth between different strain backgrounds bacterial growth was measured with the following settings: wavelength 600 nm, number of flashes 10.

### Pyochelin and pyochelin methyl ester extractions from agar plates

*P. aeruginosa* and *S. aureus* cells were grown on agar plates as described for MALDI MSI. Agar pieces containing either *P. aeruginosa* or *S. aureus* colonies from single incubations or co-incubations were excised from agar plates. Agar pieces from 5 plates were pooled and extracted with 5 ml of acidified EtOAc. 2 ml of the acidified EtOAc phase were removed, centrifuged at 5,000 rpm (Biofuge pico, Heraeus) to remove nonsoluble particulates and evaporated to dryness in a SpeedVac vacuum concentrator (SPD131DDA SpeedVac Concentrator; Thermo Scientific) and stored at – 20 ° C.

### Pyochelin and pyochelin methylester extractions from liquid cultures

Ethyl acetate (EtOAc; J.T. Baker and Fisher Scientific) and Optima-grade methanol (MeOH; Fisher Scientific) were used. *P. aeruginosa* PAO1 wild type or Δ*pqsA* cells were grown overnight in LB (RPI) at 37 °C with 200 rpm shaking. Bacterial cells were centrifuged at 5,000 rpm (Biofuge pico, Heraeus) and washed with 0.5 % chelex treated CAA. New cultures were seeded in 0.5 % chelex treated CAA at a starting OD_600_ of 0.05 and incubated at 37 °C with 200 rpm shaking. After 48 h the cultures were centrifuged at 3,800 rpm (Eppendorf Centrifuge 5810 R) for 30 min at room temperature. The supernatant was decanted and filtered using a Nalgene Rapid-flow Filter Unit (0.2 µm aPES membrane; Thermo Scientific). The cell free supernatant was acidified to pH 1.8 – 2 with 6 M HCl. Pyochelin and pyochelin methylester were extracted with 3 volumes EtOAc and evaporated to dryness in a Rotavapor (Büchi; Rotavapor R-300) or a SpeedVac vacuum concentrator (SPD131DDA SpeedVac Concentrator; Thermo Scientific). Dried samples were resuspended in MeOH and stored at – 20 ° C. Samples were centrifuged for 5 min at 10,000 rpm (Thermo; Sorvall ST 40R) to remove nonsoluble particulates and diluted as needed in MeOH containing 1 µM glycocholic acid.

### Micro plug solvent screen

Extraction of the unknown metabolite at *m/z* 339 was tested with 15 different solvents (**Table S3**). Briefly, *P. aeruginosa* Δ*pqsA* and *S. aureus* cells were grown in co-cultures on agar plates as described for MALDI-TOF MSI. Agar plugs from the interaction zone (highest intensity of *m/z* 339 in MALDI-TOF MSI) were harvested with the bottom end of a 100 µl pipette tip. For each solvent, plugs from 4 interaction plates were pooled, homogenized with a pipette tip and incubated with 200 µl solvent for 1 h at room temperature. Tubes were centrifuged at 5,000 rpm (Biofuge pico, Heraeus), 150 µl of clean solvent were transferred in a fresh tube, evaporated to dryness in a SpeedVac vacuum concentrator (SPD131DDA SpeedVac Concentrator; Thermo Scientific) and stored at – 20 ° C.

### LC-MS guided transposon mutant screen

*P. aeruginosa* Δ*pqsA* was grown in 50 ml of 0.5% chelex treated CAA supplemented with 0.4 mM MgCl_2_ at pH 7.4 for 48h at 37 °C with shaking. The *P. aeruginosa* culture was centrifuged at 3,800 rpm in an Eppendorf Centrifuge 5810 R at room temperature for 20 min. The supernatant was filtered with a Nalgene Rapid-Flow Filter Unit (0.2 µm aPES membrane; Thermo Scientific). The cell free supernatant was used for co-incubation with the *S. aureus* transposon mutants.

*S. aureus* transposon mutants were grown in 96-well plates in 220 µl TSB at 700 rpm at 37°C (Microtiter Plate Shaker Incubator SI505; Stuart) for 18 h. 200 µl of each grown culture were transferred to a 96-deep well plate, mixed with 400 µl cell free *P. aeruginosa* Δ*pqsA* supernatant. The plates were sealed with Gas permeable sealing membranes (Breathe-Easy; Diversified Biotech; 9123-6100) and incubated at 37 °C with 800 rpm shaking at 37 °C (Microtiter Plate Shaker Incubator SI505; Stuart). *S. aureus* train JE2, the parental strain of the transposon library was used as a control. After incubation, proteins were crashed out by addition of 300 µl ethyl acetate and all samples were evaporated to dryness in a SpeedVac vacuum concentrator (SPD131DDA SpeedVac Concentrator; Thermo Scientific). Dried samples were stored at −20C until further processing. Samples were resuspended in 200 µl Optima-grade MeOH and incubated in a sonicator bath (Branson; 5800) at room temperature for 40 min and an additional 30 min in the dark at room temperature. Plates were centrifuged at 10,000 rpm (Biofuge pico, Heraeus) and clear MeOH extracts were diluted 1:10 v/v with MeOH containing 1 µM glycocholic acid for subsequent analysis by LC-MS.

### Murine model of skin infection

*P. aeruginosa* strain PAO1 was grown overnight in 5 mL of LB medium at 37°C shaking at 200 rpm. *S. aureus* strains of LAC USA300 (MRSA WT, Δ*spm* and Δ*spm*::*spm*) were all grown overnight in 5mL of TSB medium at 37°C shaking at 200 rpm. For mouse infections, overnight cultures of MRSA and PAO1 were diluted 1:100 into 35 mL TSB and LB, respectively, and grown shaking in flasks at 200 rpm to an OD_600_ of 0.5 (≈2 hr for MRSA and ≈3 hr for PAO1). Subcultured bacteria were then pelleted and resuspended in sterile saline, so that 10 µL of each culture was normalized to ≈5x10^5^ CFU. 1 mL of each strain suspension was aliquoted into 1.5 mL Eppendorf tubes and kept on ice for murine infection inocula.

All *in vivo* infections were on female C57BL/6 WT mice obtained from Jackson Laboratories (Bar Harbor, ME). All animals were housed and maintained at the University of Colorado Anschutz Medical Campus Animal Care Facility accredited by the Association for Assessment and Accreditation of Laboratory Care International (AAALAC) and mice were allowed to acclimate to the BSL-2 level facility for at least seven days prior to their inclusion in this study’s in vivo murine model. All animal studies described herein were performed in accordance with best practices outlined by the Office of Laboratory Animal Resources (OLAR) and Institutional Animal Care and Use Committee (IACUC) at the University of Colorado (protocol #00987).

A murine model of wound infection was used to assess chemical interactions between MRSA and PAO1 during polymicrobial infection. One day prior to infection, mice were anesthetized (inhalation of isoflurane, 2-3%), and the fur on the dorsal surface was carefully shaved and Nair applied to completely expose the skin. On day 0, mice were anesthetized, and exposed dorsal skin was sterilized with a PVP iodine prep pad (PDI Healthcare, Woodcliff Lake, NJ). Bupivacaine hydrochloride was injected subcutaneously with a 30-gauge insulin syringe at a dose of 1-2 mg/kg in the medial thoracic region as an analgesic for the area to be wounded. A disposable 6 mm biopsy punch (Integra Miltex) was used alongside dissection scissors and forceps to excise a circular section of skin 6 mm in diameter to create a wound. For single infections, 10 µL (5×10^5^ CFU) of MRSA inoculum in sterile saline was inoculated into the open wound by pipette. For co-infections, 5 µL (2.5×10^5^ CFU) of both MRSA and PAO1 were inoculated into the open wound, for a final inoculating dose of 5×10^5^ CFU.

Following inoculation, the wounds were covered with the transparent dressing Tegaderm, to allow for observation and prevent contamination, followed by 2 bandages to protect the wounded area. Groups of at least 5 mice were used for each test condition per experiment. Animals were euthanized by CO_2_ inhalation followed by cervical dislocation as a secondary method of euthanasia at the completion of the experiment. However, some mice were sacrificed early to minimize distress and prevent spontaneous mortality according to the University of Colorado Guidelines for Establishing Humane Endpoints in Animal Study Proposals. Each animal’s physical condition was monitored daily to evaluate signs of morbidity (including body weight, physical appearance, clinical signs, unprovoked behavior, or response to external stimuli) that may consititute an early endpoint as the most humane option. Therefore, if animals exhibited signs of distress according to these guidelines, such as weakness, inability to drink water or eat, signs of systemic infection, or severe weight loss, mice were euthanized.

At daily timepoints over the course of 3 days, animals were monitored, weights measured, and wounds photographed adjacent to a ruler. Baseline body weights of mice were measured before infection, and everyday thereafter for 3 days or until animals were sacrificed before the third day of infection using a laboratory scale. Wound sizes were analyzed from photos taken daily with a Canon Rebel Powershot, and measured by scaling and determining wound area using ImageJ software (National Institutes of Health, NIH) calculated using the following equation: (A_0_-A_t_)/A_0_•100.

### Pyochelin and pyochelin methyl ester extractions from mouse tissue

Homogenized mouse skin tissue was acidified with 3 drops of 6 M HCl and transferred to a scintillation vial. To remove remaining homogenate, tubes were washed twice with 500 µl EtOAc, the solvent was transferred and pooled with the homogenate into the same scintillation vial. An additional 1 ml of EtOAc was added to the homogenate and the samples were mixed by pipetting. Phases were allowed to separate for 5 min and 1 ml of the EtOAc phase was transferred to a fresh tube. Homogenate was extracted three times, yielding a total of 3 ml of the EtOAc phase, which was concentrated to dryness in a SpeedVac vacuum concentrator (SPD131DDA SpeedVac Concentrator; Thermo Scientific). Samples were stored at – 20 ° C until further use.

To prepare mouse skin tissue extracts for LC-MS analysis, samples were thawed on room temperature, resuspended in 120 µl MeOH, incubated for 40 min in a sonicator bath (5800; Branson) at room temperature and an additional 30 min in the dark at room temperature. Samples were mixed by pipetting, centrifuged for 2 min at 10,000 rpm (Sorvall ST 40R; Thermo) to remove nonsoluble particulates and 100 µl of clean extract were transferred to a fresh tube. Samples were stored at – 20 ° C or directly diluted 1:10 (v/v) in MeOH containing 1 µM glycocholic acid.

The tissues were then homogenized in three 30 second increments (for a total of 90 seconds) using a bead beater, serially diluted, and selectively plated for CFU counts. All tissue samples were plated on mannitol salt agar (MSA) with cefoxitin to select for MRSA and *Pseudomonas* Isolation Agar to select for PAO1. Tissue samples were then frozen at − 80°C to preserve for later chemical analysis.

### MALDI-TOF MSI

Overnight cultures of MRSA or PAO1 (WT, Δ*pqsA* or Δ*pqsA* Δ*pch*) were diluted to OD_600_ of 0.2 and 5 µl of the bacterial solution were spotted on tryptic soy agar plates (TSA; Remel) supplemented with agar (2% (w/v) final concentration). For bacterial co-incubations, the bacterial solutions were spotted 5 mm apart. Samples were prepared for MALDI-MSI as previously described^40,41^. Briefly, for each sample a region of agar including the bacterial colonies was excised from the culture, laid on top of a MALDI MSP 96 anchor plate (Bruker Daltonics). Universal matrix (1:1 2,5-dihydroxybenzoic acid (DHB): α-cyano-4- hydroxycinnamic acid (CHCA); (Sigma-Aldrich)) was applied manually using a 53 µm molecular sieve. Samples were dried at 37°C overnight and a photograph was taken. All colonies were subjected to MALDI-TOF MSI in positive reflectron mode using 500 µm spatial resolution in both X and Y dimensions by a Bruker Daltonics Autoflex Speed. Two dimensional metabolite distribution was visualized using Bruker FlexImaging version 4.0. False colored images were optimized for visualization of ion distribution.

### Metabolomics data collection

Mass spectrometry data acquisition was performed using a Bruker Daltonics Maxis II HD quadrupole time of flight (qTOF) mass spectrometer with a standard electrospray ionization (ESI) source. Mass spectrometer tuning was performed by infusion with tuning mix ESI-TOF (Agilent Technologies) at a 3-μl/min flow rate. As a lock mass internal calibrant, hexakis (1H,1H,2H-difluoroethoxy)phosphazene ions (Apollo Scientific, *m*/*z* 622.1978) located on a wick within the source was used. Samples were introduced by an Agilent 1290 ultraperformance liquid chromatography (UPLC) system using a 10-μl injection volume. The solvent system for UPLC separation consisted of Optima-grade water (Fisher Scientific) and acetonitrile (ACN, Fisher Scientific) (Buffer A: 98:2; Buffer B: 2:98) with 0.1% Formic acid (FA, Fisher Scientific). Extracts were separated using a Phenomenex Kinetex 2.6-μm C18 column (2.1 mm by 50 mm) or a Phenomenex Kinetex 1.7-μm C18 column (2.1 mm by 50 mm) using a 9 min, linear water-ACN gradient at a flow rate of 0.5 ml/min. To compare the retention times of isolated and synthetic Pch and MetPch isomers, samples were separated using a Phenomenex Luna 5 µm C18(2) column (250 mm by 2 mm) using a 20 min, linear water-ACN gradient at a flow rate of 0.5 ml/min. The mass spectrometer was operated in data-dependent positive ion mode, automatically switching between full-scan MS and MS/MS acquisitions. Full-scan MS spectra (*m/z* 50 to 1,500) were acquired in the TOF-MS, and the top five most intense ions in a particular scan were fragmented via collision-induced dissociation (CID) using the stepping function in the collision cell.

### Molecular networking

A classical molecular network was created from the mzXML files using the online workflow (version release 30) (https://ccms-ucsd.github.io/GNPSDocumentation/) on the GNPS website (http://gnps.ucsd.edu)46. Briefly, the data was filtered by removing all MS/MS fragment ions within +/− 17 Da of the precursor m/z. MS/MS spectra were window filtered by choosing only the top 6 fragment ions in the +/− 50Da window throughout the spectrum. The precursor ion mass tolerance was set to 0.05 Da and a MS/MS fragment ion tolerance of 0.1 Da. A network was then created where edges were filtered to have a cosine score above 0.7 and more than 6 matched peaks. Further, edges between two nodes were kept in the network if and only if each of the nodes appeared in each other’s respective top 10 most similar nodes. Finally, the maximum size of a molecular family was set to 100, and the lowest scoring edges were removed from molecular families until the molecular family size was below this threshold. The spectra in the network were then searched against GNPS’ spectral libraries. The library spectra were filtered in the same manner as the input data. All matches kept between network spectra and library spectra were required to have a score above 0.7 and at least 6 matched peaks. The molecular network (https://gnps.ucsd.edu/ProteoSAFe/status.jsp?task=b225a233163545b7b8eba55c53fff754) was visualized in Cytoscape (version 3.8.2)^60^.

### Total synthesis of pyochelin and methyl ester pyochelin

All reactions were carried out in flame dried glassware under argon unless water was used in the reaction. Reagents and anhydrous solvents were purchased from Sigma-Aldrich, Acros Organics or TCI and used without further purification unless otherwise stated. Reactions were monitored using silica gel 60 F254 TLC plates. Plates were visualized with 254 nm UV light or when appropriate a basic KMnO_4_ solution or 5% solution of FeCl_3_ in methanol. Flash chromatography was performed using 60 μm mesh standard grade silica gel from Millipore Sigma. NMR solvents were obtained from Cambridge Isotope Labs and used as is. All ^1^H NMR (400 MHz) were recorded at 25 °C on a Bruker Avance spectrometer. All ^13^C NMR (101 MHz) spectra were also recorded at 25 °C on Bruker Avance spectrometers. Chemical shifts (δ) are given in parts per million (ppm) relative to the respective NMR solvent; coupling constants (J) are in hertz (Hz). Abbreviations used are s, singlet; d, doublet; dd, doublet of doublets; td, triplet of doublets; m, multiplet. All high-resolution mass spectrometry measurements were made in the Mass Spectrometry and Proteomics Facility at the University of Notre Dame. Preparatory HPLC purification was performed on a Preparatory HPLC purification was performed on an Agilent 1260 Infinity HPLC fitted with a Jupiter 4 μm Proteo 90 Å (250 x 21.2 mm) LC column running a 45 minute method with a gradient of 20- 95% MeCN in Water with 0.1% TFA running a gradient of 20-95% MeCN in Water with 0.1% TFA.

#### (S)-2-(2-hydroxyphenyl)-N-methoxy-N-methyl-4,5-dihydrothiazole-4-carboxamide

To a solution of 2-hydroxybenzonitrile 8 (2.5 g, 21 mmol) in methanol (93 mL) was added D-Cysteine HCl hydrate (3.7 g, 21 mmol) and phosphate buffer 0.1 M (pH 6.4) (75 mL). The solution was adjusted to pH 6.4 by addition of solid K_2_CO_3_ and the reaction mixture was stirred at 60 °C overnight. The mixture was concentrated under reduced pressure, and the yellow crude material was diluted with water (100 mL). The solution was adjusted to pH 2.0 by addition of solid citric acid. After extraction with DCM (3 x 100 mL), the organic layers were collected, dried over Na_2_SO_4_ and filtered. The solvent was removed under reduced pressure to afford the as a yellow powder (4.2 g, 90%).

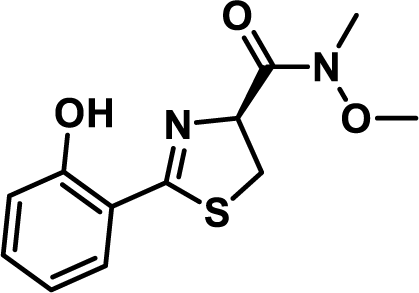

To a flamed dried reaction flask charged with carboxylic acid (3.8 g, 17 mmol) under argon was added anhydrous DMF (95 mL). Then, EDC (13 g, 68 mmol), HOBt (13 g, 68 mmol), N,O-dimethyl hydroxylamine hydrochloride (3.3 g, 34 mmol) were added. Triethylamine (9.5 mL, 68 mmol) was added dropwise by syringe. The reaction mixture was stirred overnight at room temperature. The reaction was then diluted with water and the mixture was brought to pH 2 by addition of 1 M HCl and extracted with ethyl acetate (3 x 100 mL). The combined organic layers were washed with brine, dried over anhydrous sodium sulfate, filtered, and concentrated. The crude material was purified by silica gel chromatography (10% to 100% EtOAc in hexanes) to give an off-white solid (2.5 g, 55% yield). NMR values matched previously reported spectra ^61^.

#### N-methyl-L-cysteine hydrochloride

A solution of L-cysteine hydrochloride (16.0 g, 102 mmol) in water (10 mL) and NaOH (1 M, 10.2 mL, 10.2 mmol) was added aqueous formaldehyde (11.3 mL, 152 mmol, 37% by weight). After stirring at room temperature for 24 hours, the reaction was cooled to 0 °C and ethanol (25 mL) and pyridine (12 mL) were added. The resulting white precipitate was filtered and dried under vacuum (12.5 g, 93%). Used in next step without further purification.

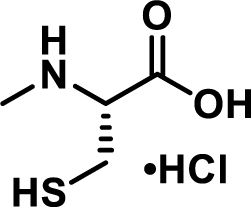

The carboxylic acid (5.0 g, 38 mmol) from the previous step was dissolved in liquid ammonia (100 mL) condensed at −78 °C. Water (0.68 mL, 38 mmol) was added. Sodium metal was added until the solution retained a deep blue color. The ammonia was allowed to evaporate, and the reaction was quenched with sat. ammonium chloride. After drying, the residue was dissolved in water and acidified to pH 1 with 6N HCl and water was removed under vacuum. The remaining residue was washed multiple times with hot ethanol and the organic layer was concentrated under vacuum to afford the product as a sticky white solid (2.4 g, 37%). NMR values matched previously reported spectra ^62^. DMF was used as internal standard to calculate purity of amino acid by weight to ensure no inorganic salts were present.

#### Pyochelin I and II

Weinreb amide 13 (2.5 g, 9.4 mmol) was added to a flame-dried flask under argon and dissolved in dry THF (190 mL). The reaction mixture was brought to −78 °C in a bath of dry ice and acetone. Lithium aluminum hydride (4M, 2.6 mL, 10 mmol) was added dropwise by syringe down the side of the reaction flask into the reaction mixture. The reaction was stirred at between −50 and −30 °C and monitored by TLC. After 2 hours, the reaction was quenched by slow addition of saturated NH_4_Cl and the reaction was allowed to warm to room temperature while stirring vigorously until two phases formed. The phases were separated, and the aqueous layer was extracted with EtOAc (3 x 25 mL). The combined organic layers were washed with brine, dried over anhydrous sodium sulfate, filtered, and concentrated. Due to instability the aldehyde was used as quickly as possible in the cyclization without any purification.

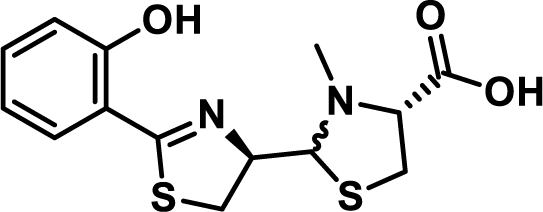

Potassium acetate (4.1 g, 42 mmol) and N-methyl-L-cysteine hydrochloride (2.4 g, 14 mmol) were added to a solution of the crude aldehyde in EtOH (250 mL) and H2O (62.5 mL). The mixture was stirred at 25 °C overnight and then diluted with H_2_0, adjusted to pH 4.5 with 6 N HCl then extracted with EtOAc. The organic layers were worked up to yield crude pyochelin as a mixture of 4 isomers. HPLC (15 cm, C12 column) analysis showed the four separate isomers eluting at 13.65, 14.82, 15.96 and 17.12 minutes on a 30 min 5-95 % ACN (0.1% FA) LC method (LC trace below). Each of the four major peaks has the correct mass of [M+H]^+^ 325 m/z. The natural pyochelin isomers were determined to be the second and third peaks on LC based on the ^1^H NMR shifts of their methyl esters. Peaks one and four are neopyochelin. Purification of 125 mg of crude material on prep HPLC (20-95 ACN in aq. 0.1% TFA) led to 40 mg each of pyochelin and neopyochelin as their bis-trifluoro acetate salts. 15 mg of each was put aside then the rest was dissolved in a pH 4.5 phosphate buffer (680 mg potassium dihyrdrogen phosphate in 100 mL DI water) then extracted with ethyl acetate and dried under high vacuum to afford a mixture of pyochelin I and II which could not be purified further due to their rapid interconversion.

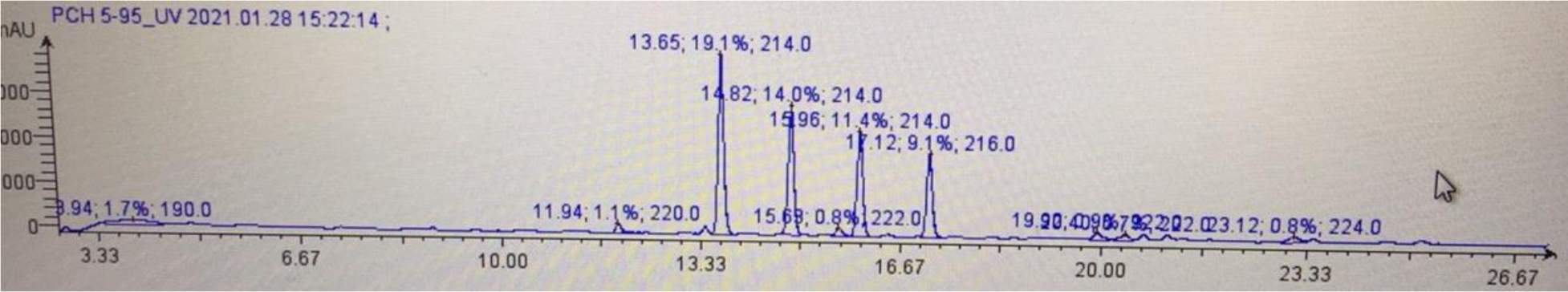
Copy of LC Trace (30 min, 5-95% MeCN in 0.1% Formic Acid)

#### Pyochelin Methylesters

To a mixture of the free bases of pyochelin I and II (100 mg, 0.3 mmol) in benzene (5 ml) and MeOH (1.25 mL) was added TMSCHN_2_ (2M, 200 uL, 0.4 mmol) at room temperature. The reaction was allowed to stir for 30 minutes before being concentrated under vacuum. The residue was then purified via prep HPLC (20-95 ACN in aq. 0.1% TFA) to afford the pyochelin methyl esters in a 4:1 ratio as their bis-trifluoro acetate salts. NMR values for pyochelin I methyl ester and pyochelin II methyl ester matched those previously reported (**Figure S5**) ^63^

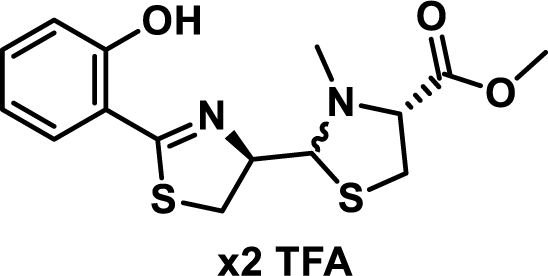

### CAS assay

CAS reagent was prepared as follows: 25 ml of 2 mM CAS reagent (Chem-Impex International) in water were mixed with 5 ml of 1 mM FeCl_3_ in 10 mM HCl, giving CAS reagent stock solution. For a 10 ml stock solution of the final CAS solution, 0.6 ml of CAS reagent stock solution were mixed with 0.4 ml of a 5 mM HDTMA solution in water and 9 ml of a 111 mM PIPES solution in water (pH 6.8). 4 mM stock solutions of synthetic pyochelin and pyochelin methylester in MeOH were serially diluted 1:2 (v/v) with MeOH, giving stock solutions of pyochelin and pyochelin methylester at 4 mM, 2 mM 1 mM, 0.5 mM, 0.25 mM and 0.125 mM concentrations. 5 µl of each stock solution were mixed with 195 µl of the final CAS solution in 96-black well plates (Corning Incorporated) and incubated at room temperate in the dark. CAS solution incubated with MeOH was used as reference. After 5 h of incubation, the fluorescence of the samples was recorded on an Infinite M PLEX plate reader (Tecan) with the following settings: Excitation 425 nm, emission 630 nm, Bandwith 9 nm, Number of flashes 25. Siderophore activity of Pch and MetPch was calculated according to the following formula: (Ar-As)/Ar x 100 % (Ar = absorbance of reference and As = absorbance of sample)

### ROS assay

Stock solutions of 2’,7’-dichlorofluorescin diacetate (DCF-DA; Sigma) were prepared by dissolving DCF-DA at a final concentration of 5 mM in absolute ethanol (Fisher Scientific) and stored at – 20 °C. The DCF-DA stock solution was then hydrolyzed to the non-fluorescent 2′,7′-dichlorodihydrofluorescein (DCF-H) by mixing 0.5 ml DCF-DA (5 mM) with 2 ml of 0.1 N NaOH at room temperate for 30 min. The reaction was stopped by adding 7.5 ml of 10x PBS pH 7.4 (0.1 M, without calcium and magnesium, Gibco). *S. aureus* cultures grown in TSB overnight were used to inoculate fresh TSB cultures at a starting OD_600_ of 0.05 and incubated at 37 °C with shaking (200 rpm). After 2.5 h, the OD_600_ of the *S. aureus* cultures was measured and all cultures were set to an OD_600_ of 1. Cultures were incubated with either Pch or MetPch at final concentrations of 10 µM, 20 µM and 40 µM or an equal volume of methanol (control) for 90 min at 37 °C with 800 rpm shaking (Microtiter Plate Shaker Incubator SI505; Stuart). Bacterial cultures containing either Pch, MetPch or methanol were adjusted to OD_600_ of 1 and centrifuged at 5,000 g (Biofuge pico, Heraeus) for 10 min. Bacterial cell pellets were resuspended in 100 µl DCF-H (50 µM) solution and incubated for 40 min. 100 µl were added to a 96-well black plate (Corning Incorporated). DCF-H incubated with sterile medium was used as control. ROS production was determined as a function of oxidation of the non-fluorescent DCF-H to 2′,7′-dichloroofluorescein (DCF) in an Infinite M PLEX plate reader (Tecan) with the following settings: Excitation 488 nm (bandwith 9 nm); emission 515 nm (bandwith 20 nm).

### *P. aeruginosa* growth support assay

*P. aeruginosa* strains were pre-grown for 36 h in M9 minimal medium supplemented with 0.5% (v/v) glycerol as carbon source, spun down and resuspended in either M9 glycerol or M9 glycerol containing 500 µM 2,2’-Bipyridine (Fisher Scientific). The bacterial stock solutions were used to inoculate M9 glycerol and M9 glycerol medium containing 500 µM 2,2’- Bipyridine at a starting OD_600_ of 0.05. Bacterial growth was followed over the course of 96 h in the presence of three different concentrations (0.25 µM, 1 µM and 2 µM) of pyochelin or pyochelin methylester or an equivalent volume of methanol. OD_600_ measurements were taken at 0 h and every 12 h thereafter in an Infinite M PLEX plate reader (Tecan).

### *S. aureus* growth inhibition assay

*S. aureus* LAC wild type was grown overnight in chelex treated TSB. Cultures were centrifuged and resuspended in chelex treated TSB. The bacterial stock solutions were used to inoculate chelex treated TSB medium at a starting OD_600_ of 0.05 and 198 µl of bacterial cultures were transferred to 96-well plates. 4 mM stock solutions of synthetic pyochelin and pyochelin methylester in MeOH were diluted to yield working solutions of 1.92 mM, 0.96 mM, 0.48 mM and 0.24 mM. 198 µl of *S. aureus* cultures at OD_600_ of 0.05 were incubated with 2 µl of Pch and MetPch working solutions to yield final concentrations of 19.2 µM, 9.6 µM, 4.8 µM and 2.4 µM of synthetic pyochelin or pyochelin methylester. 2 µl MeOh was used for control cultures. The bacterial cultures were incubated at 37 ° C with 1000 rpm shaking (Microtiter Plate Shaker Incubator SI505; Stuart) for 24 h. Bacterial growth was determined by OD_600_ measurements with an Infinite M PLEX plate reader (Tecan).

### Statistical Analysis

Statistical analyses were performed with GraphPad Prism version 9.4.1. Statistical tests performed and the P values of datasets are indicated for each individual experiment in the corresponding figure legend.

## Author Contributions

C.J., V.V.P. and A.R.H. conceptualized the study and designed experiments. C.J., K.K., J.J., M.Z. and K.S. performed experiments. C.J., K.K., M.Z., K.S. and V.V.P. analyzed data. C.J. wrote the manuscript draft. K.K., M.Z., K.S., M.S., C.M., V.V.P. and A.R.H. revised the manuscript. V.V.P. and A.R.H. supervised the study. All authors have approved the submitted version of the study and their contributions.

## Data and Materials Availability

All mass spectrometry data, including the raw files, mzXML files, metadata tables and MZmine settings are available via MassIVE (MSV000089230; *P. aeruginosa*/*S. aureus* wt, *spm* mutant, *spm* complementation interaction, MSV000089231; *P. aeruginosa*/*S. aureus* wt, *fhuG* mutant, *fhuD*1/*D*2 double mutant interaction, MSV000089233, Pyochelin methylation by *S. aureus* transposon library mutants, MSV000089234; Pyochelin methylester quantification murine skin wound model *S. aureus*/*P. aeruginosa* co-infection). Password for all datasets: MetPch2022

## Supplementary Data 1

LC-MS guided screen of Transposon Mutant library Plates A - F

## Acknowledgments

We thank Prof. Masanori Toyofuku, University of Tsukuba, for sharing *Pseudomonas aeruginosa* strains and Dr. Heidi Crosby for strain *S. aureus* LAC *fur*::tet.

