## Supplementary Material for "Pyochelin biotransformation by *Staphylococcus aureus* shapes bacterial competition with *Pseudomonas aeruginosa* in polymicrobial infections"

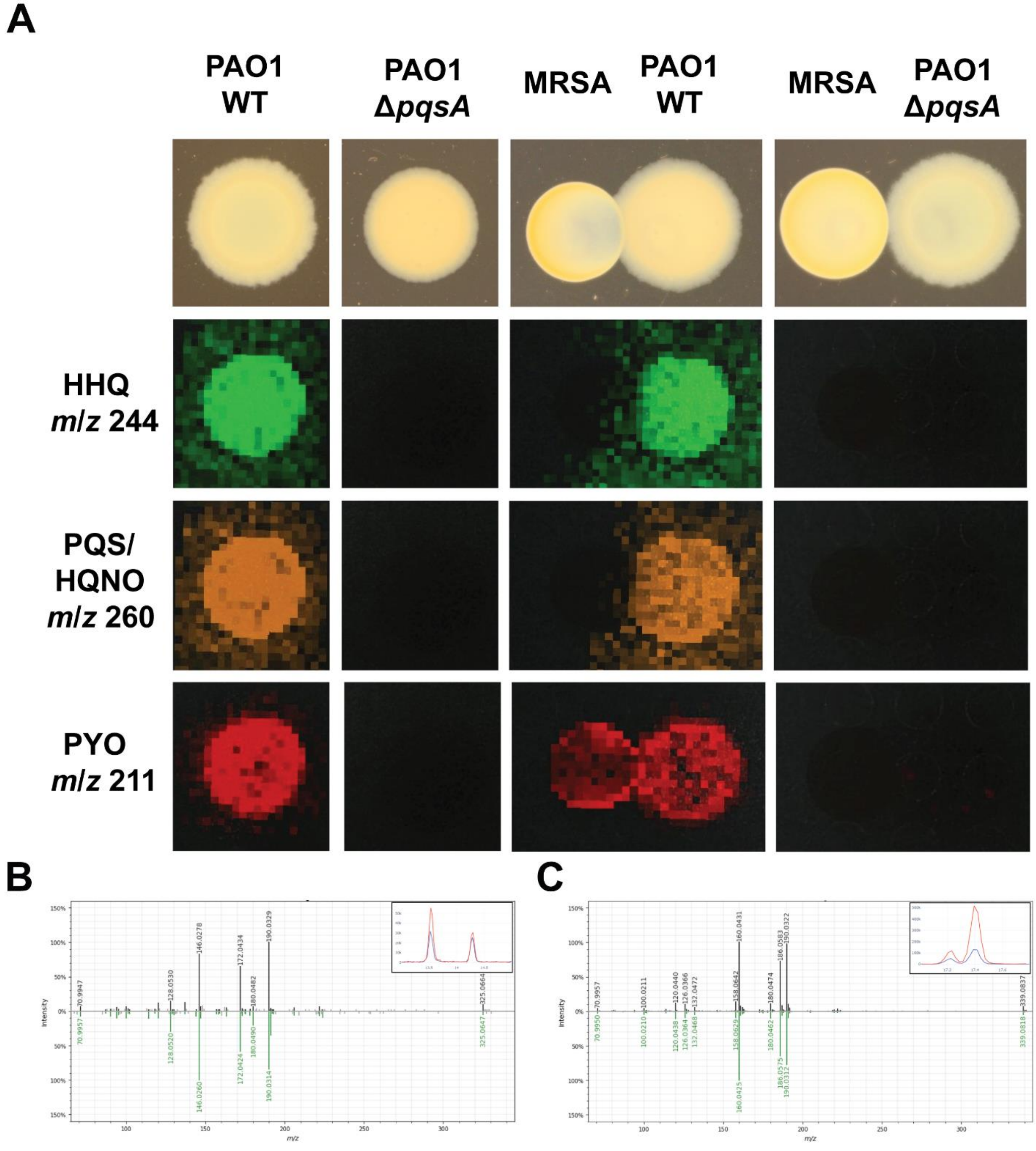

**Figure S1**: **A**) MALDI-MSI of *S. aureus* and *P. aeruginosa* grown as monocultures or interactions. Photographs of the cultures are shown on top. The second to fourth rows show the false colored *m/z* distribution for HHQ (*m/z* 244), PQS/HQNO (*m/z* 260) and pyocyanin (PYO; *m/z* 211). *P. aeruginosa* wildtype (PAO1 WT) but not a *P. aeruginosa*

quinolone mutant (PAO1 Δ*pqsA*) produces the alkyl-quinolones HHQ and PQS/HQNO as well as the quinolone regulated metabolite PYO. **B and C)** Mirror plots comparing the $MS^2$ spectrum of isolated (top; black trace) and synthetic (bottom; green trace) **B)** pyochelin and **C)** pyochelin methyl ester. Chromatogram traces of isolated (blue) and synthetic (red) metabolites are shown as inlays.

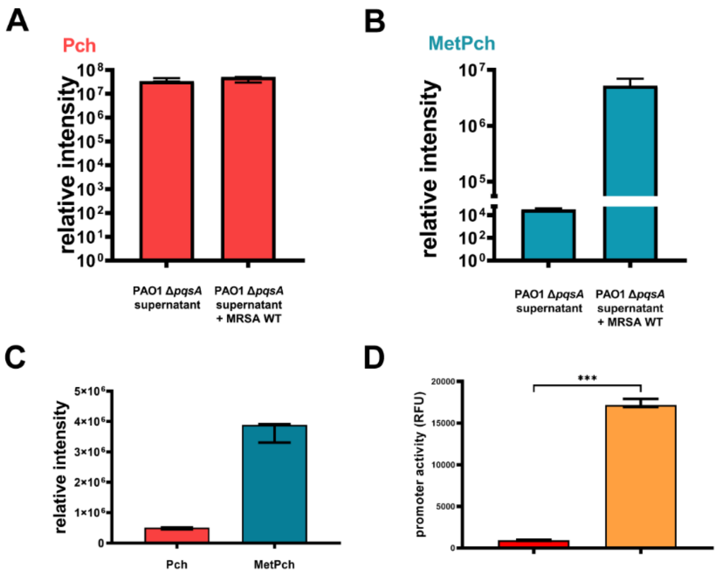

**Figure S2:** Quantification of **A)** Pch and **B)** MetPch in PAO1 Δ*pqsA* cell free SN after incubation with TSB (PAO1 Δ*pqsA* supernatant) or MRSA overnight culture (PAO1 Δ*pqsA* supernatant + MRSA WT) for 8h. **C)** Equimolar amounts of synthetic Pch and MetPch quantified by LC-MS. MetPch shows a 10-fold higher singal intensity than Pch, most likely due to more efficient ionization. **D)** The FhuD1/Smt promoter activity is elevated in a *fur* mutant (*fur*::*tet*) compared to its parent strain (WT). n = 8 biological replicates. Values are mean ± SDs ****P* ≤ 0.001 [Mann Whitney U test].

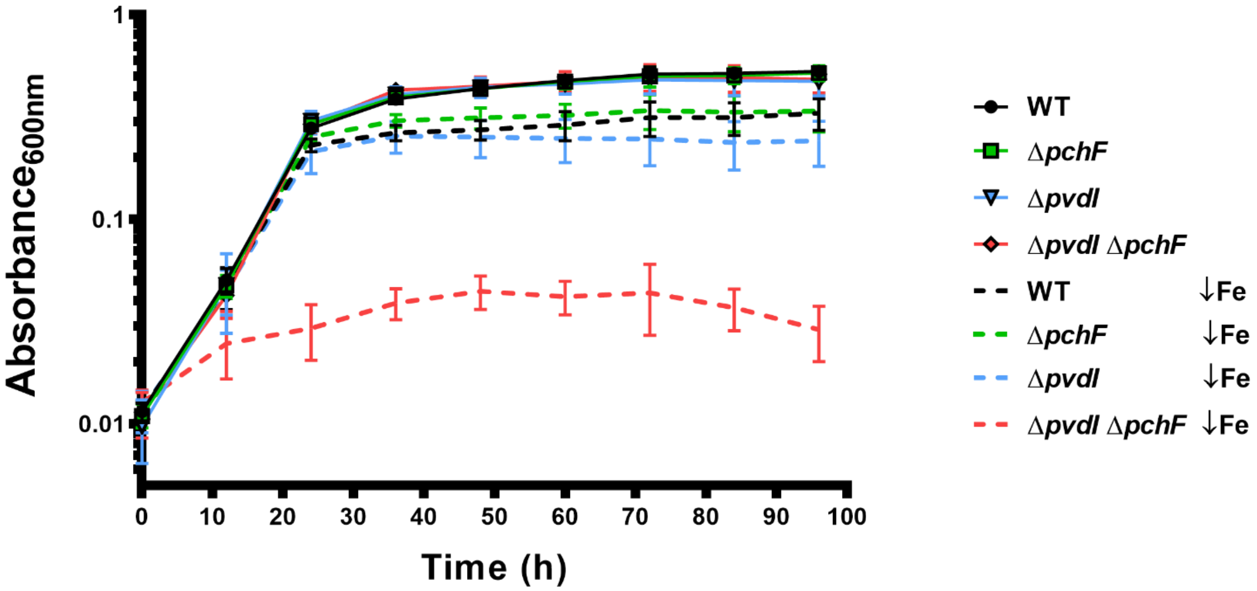

**Figure S3**: Siderophore biosynthesis is essential for growth under iron limited conditions. 96-hour growth curves of *P. aeruginosa* PAO1 wild type (WT), a pyochelin mutant (Δ*pchF*), a pyoverdine mutant (Δ*pvdI*) and a mutant deficient in production of both siderophores (Δ*pchF* Δ*pvdI*). Cultures were grown in M9 minimal medium with glycerol as carbon source. To create a low iron environment (↓Fe), 500 μM 2,2'-Bipyridine was added to the growth medium. Growth curves of bacteria grown in standard M9 medium are shown as full lines and growth curves from low iron environment are shown as dashed lines. n = 5 biological replicates from two independent experiments.

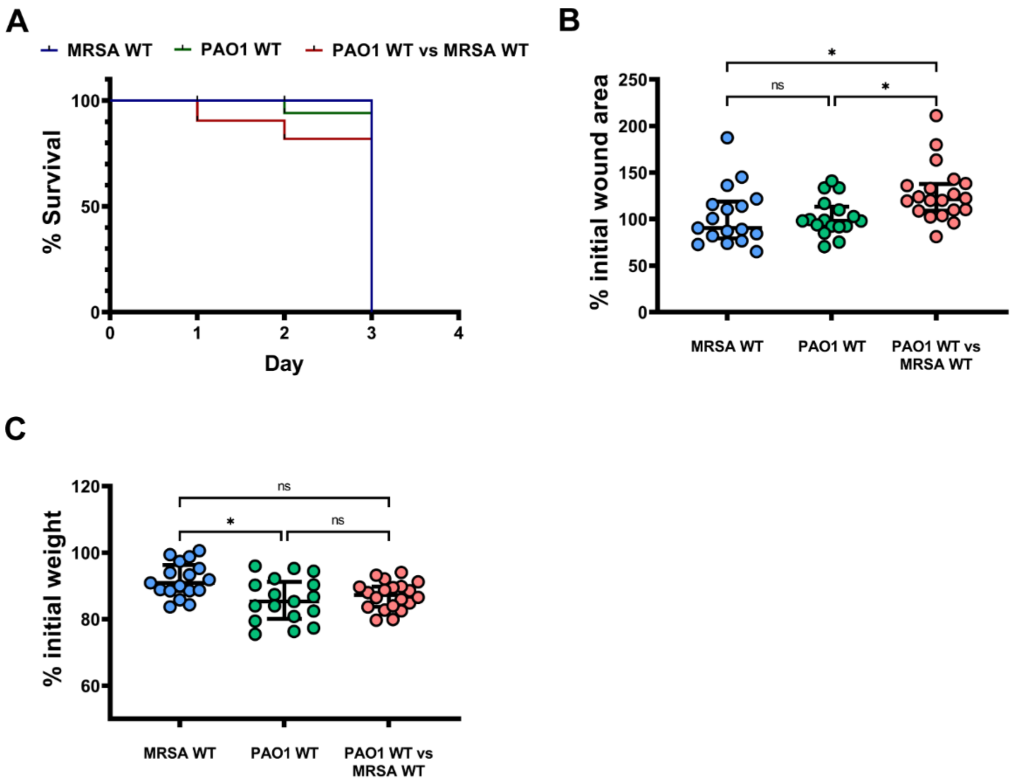

**Figure S4**: Polymicrobial infection leads to increased mortality and wound size progression. **A**). Murine survival after skin wound infection with either *S. aureus* LAC wild type (MRSA WT) or *P. aeruginosa* PAO1 wild type (PAO1 WT) or co-infection (PAO1 WT vs MRSA WT). Co-infections lead to lowered murine survival compared with single infections. **B**) The change in murine skin wound size area between start and end of the experiment. Changes were calculated from the measured wound size area before bacterial infection and at the endpoint of the murine skin wound model. The changes are expressed as percent (%) of the initial wound size area at the endpoint of the experiment. **C**) The change in murine weight between start and end of the experiment. Changes were calculated from the murine weight before bacterial infection and at the endpoint of the murine skin wound model. The changes are expressed as percent (%) of the initial murine weight at the endpoint of the experiment. **B – C**) n = 17 biological replicates from 3 independent experiments. [Kruskal-Wallis with Dunn's multiple comparison]. Values are median ± interquartile range. Each dot represents values from a single mouse. *$P \leq 0.05$, **$P \leq 0.01$, ***$P \leq 0.001$, ****$P \leq 0.0001$, ns = not significant.

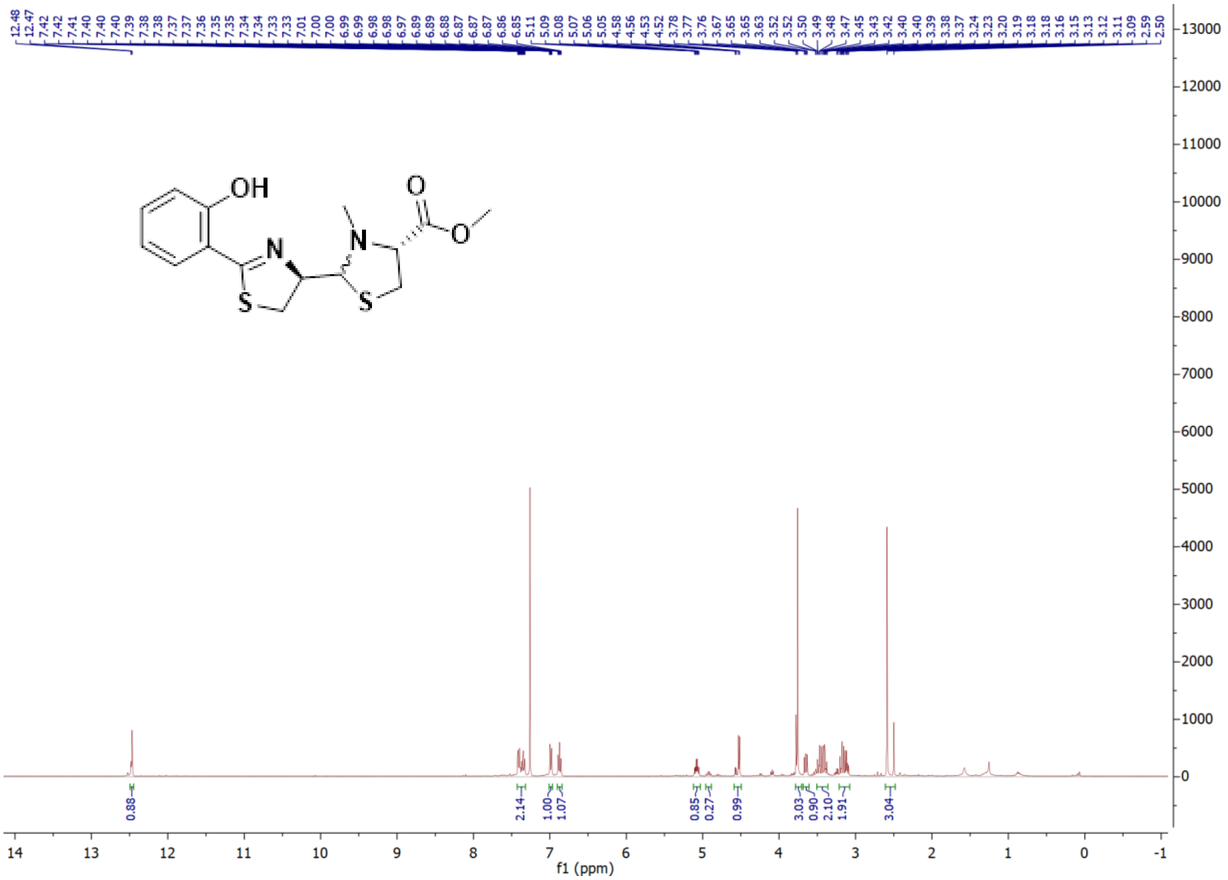

**Figure S5:** $^{1}$H NMR of MetPch (400 MHz, Chloroform-*d*) [Major Isomer] δ 12.47 (s, 1H), 7.43 – 7.32 (m, 2H), 6.99 (ddd, *J* = 8.3, 3.4, 1.2 Hz, 1H), 6.87 (td, *J* = 7.6, 1.2 Hz, 1H), 5.08 (td, *J* = 8.8, 5.1 Hz, 1H), 4.52 (d, *J* = 5.2 Hz, 1H), 3.76 (s, 3H), 3.65 (dd, *J* = 9.2, 6.3 Hz, 1H), 3.51 – 3.36 (m, 2H), 3.22 – 3.08 (m, 2H), 2.59 (s, 3H).

**Table S1:** bacterial strains and plasmids used in this study

| strain | description | source |
|---|---|---|
| ***Staphylococcus aureus*** | | |
| USA300 LAC/MRSA WT | USA300 CA-MRSA, Erm$^S$ (LAC*) | [S1] |
| USA300 LAC Δ*spm*/MRSA Δ*spm* | unmarked *spm* deletion mutant | this study |
| USA300 LAC Δ*spm*::*spm*/MRSA Δ*spm*::*spm* | repaired *spm* deletion mutant; NcoI restriction site located 110 bp downstream of *spm* ORF used as water mark | this study |
| USA 300 LAC *fur*::tet | *fur* mutant in strain LAC with tetracycline resistance | this study |
| AH2367 | *fur* mutant in strain 8325-4 with tetracycline resistance | [S2] |
| ***Pseudomonas aeruginosa*** | | |
| PAO1 | PAO1 wild type | [S3] |
| PAO1 Δ*pqsA* | unmarked *pqsA* deletion mutant; deficient in alkyl-quinolone production | [S4] |
| PAO1 Δ*pvdI* | unmarked *pvd*I deletion mutant; deficient in pyoverdine production | this study |
| PAO1 Δ*pchF* | unmarked *pchF* deletion mutant; deficient in pyochelin production | this study |
| PAO1 Δ*pvdI* Δ*pchF* | unmarked *pvdI* and *pchF* double deletion mutant; deficient in pyoverdine and pyochelin production | this study |
| PAO1 Δ*pch* | unmarked pyochelin biosynthesis cluster mutant; deficient in pyochelin production | this study |
| ***Escherichia coli*** | | |
| DC10b | Cloning strain (*dcm*-) | [S5] |
| Top10 | F- mcrAΔ(mrr-hsdRMS-mcrBC) φ80lacZΔM15 ΔlacX74 nupG recA1 araD139 Δ(ara-leu)7697 galE15 galK16 rpsL(StrR ) endA1 λ- | Invitrogen |
| **plasmids** | | |
| pJB38 | Mutation generation vector, Amp$^R$ (*E.coli*), Cm$^R$ (*S. aureus*) | [S6] |
| pEX2G | Mutation generation vector, | [S7] |
| pHC48 | *S. aureus* constitutive DsRed expression vector | [S8] |
| pJB*spm* | pJB38-*spm*; construct for unmarked deletion of *spm* | this study |
| pJB*spm*KI | pJB38-*spm*KI; construct to repair Δ*spm* mutant strain; water mark: NcoI restriction site | this study |
| pJB*fhuD*1 | pJB38-*fhuD*1; construct for unmarked deletion of *fhuD*1 | this study |
| pJB*fhuD*2 | pJB38-*fhuD*2; construct for unmarked deletion of *fhuD*2 | this study |
| pJB*fhuG* | pJB38-*fhuG*; construct for unmarked deletion of *fhuG* | this study |
| pEX*pch* | pEX2G-*pch*; construct for unmarked deletion of the entire pyochelin biosynthesis cluster | this study |
| pEX*pchF* | pEX2G-*pchF*; construct for unmarked deletion of *pchF* | this study |
| pEXpvdI | pEX2G-*pvdI*; construct for unmarked deletion of *pvdI* | this study |
| pHC*spm*prom | pHC48-*spm*prom, promoter region of the *spm*/*fhuD*1 operon with DsRed on the pHC48 backbone. | this study |

**Table S2:** primer used in this study

| primer name | primer sequence |
|---|---|
| **construction primer** | |
| spm_up_fw_KpnI | ccccggtacCGTGTTCAGCATCTGTCGAT |
| spm_up_rv_NcoI | ggggccatggTTTTGCTCGAGCTATCAAAGG |
| spm_dw_fw_NcoI | ggggccatggGTTCCAAAGAGACAGAGACATTAAG |
| spm_dw_rv_SalI | ccccgtcgacGGCTACTGCTGGCGAAGA |
| fhuD1_up_fw_KpnI | ccccggtaccAGAGGGGCACCACCAATAA |
| fhuD1_up_rv_NcoI | ggggccatggATCGACAGATGCTGAACACG |
| fhuD1_dw_fw_NcoI | ggggccatggGAAGATTATTGGTTCACAGATCCT |
| fhuD1_dw_rv_SalI | ccccgtcgacTCCCTAGAAAACAGCAGCTCA |
| fhuD2_up_fw_EcoRI | ccccgaattcAGTAAAGCGCCAAATGGTTG |
| fhuD2_up_rv_KpnI | ccccggtaccTTTCATAATTTCCTCCTATTGAAAATG |
| fhuD2_dw_fw_KpnI | ccccggtaccGCTGGTACATACTGGTACAACGA |
| fhuD2_up_fw_KpnI_new | ccccggtaccAGTAAAGCGCCAAATGGTTG |
| fhuG_up_fw_EcoRI | ccccgaattcTTTGCTGGATTTTTAGGTGCT |
| fhuG_up_rv_NcoI | ggggccatggTGCTATCAATTGTCTGCGTTT |
| fhuG_dw_fw_NcoI | ggggccatggATTGGTGCACCGTATTTCTT |
| fhuG_dw_rv_SalI | ccccgtcgaCCTTCCATTTCTGCAAGCTC |
| pchF_up_fw_XbaI | cccctctagaCGCGCCTGCTCGAAG |
| pchF_up_rv_HindIII | ccccaagcttCTCGCTCCAGAGTTCGATG |
| pchF_dwn_fw_HindIII | ccccaagcttCTGGCGCAATCCTTGTC |
| pchF_dwn_rv_EcoRI | ccccgaattCTCCAGCACCTGTTCCAG |
| pch_up_fw_EcoRI | ccccgaattCACTGCCTTCAACCTGCTC |
| pch_up_rv_NcoI | ggggccatggGTCGCCTGAAAACGAAGAC |
| pch_dwn_fw_NcoI | ggggccatggGCGGGCGAGGAAAGTT |
| pch_dwn_rv_XbaI | cccctctagaCGGTAGGTGTCCCACTTGA |
| pvdI_up_fw_XbaI | ggggtctaGATGCCGTCGTTGGTCAG |
| pvdI_up_rv_NcoI | ggggccatggACGCTTCTCAACGGGTAGTT |
| pvdI_dw_fw_NcoI | ggggccatggAGCGAACTAGAGGCGATCTG |
| pvdI_dw_rv_EcoRI | ccccgaattcTGAACATGGTCACACCCTTG |
| **colony PCR primer** | |
| fhuD2_chrom_col_fw | TGCCGATAATTGGGACGTAT |
| fhuD2_chrom_col_rv | GCATGCATACGCCATTCTTT |
| fhuG_chrom_col_fw | TTCAGGTGCTTCATTTGCTTT |
| fhuG_chrom_col_rv | ACGCTTCTACACGCGATTTT |
| spm_chrom_col | CCCCCTTTTCAACTGACAGA |

| | |
|---|---|
| fhuD1_chrom_col | CAAAAATAAACGCAACTGCAA |
| pch_chrom_col_up | CTGATCCTCCACGCCATC |
| pch_chrom_col_dwn | ATCGGCTTCCTGGTATTCG |
| pchF_chrom_col_up | CTCGACGAGGCGTTGC |
| pchF_chrom_col_dwn | CGATAGGTGCGCAGCAG |
| pvdI_chrom_col_up | TTGCCGGTATAAGGGTTCAG |
| pvdI_chrom_col_dwn | GCCTGGTGATTGAACATGAC |
| HC479 | AGAAAAGCTTGCATTTTATTGAGAA |
| HC480 | TTCAATAATTGTTTCCATAACCACA |
| **mutant complementation primer** | |
| spm_KI_up_fw_short_KpnI | ccccggtaccGGATTGAAGAGTGGGACGAT |
| spm_KI_up_rv_NcoI_NcoI_SalI | ccccgtcgacatatccatggTCCCTAGAAAACAGCAGCTCA |
| spm_KI_dw_fw_NcoI | ggggccatggTGTCCCATTCCTTTTACGAGA |
| spm_KI_dw_rv_short_SalI | ccccgtcgaCAAGCAATCACCGACCTGTA |
| **sequencing primer** | |
| fhuD2_dw_seq | GTGCCGTTGTTATCGTTCAAT |
| fhuD2_up_seq | TGCTGCATCTTCCATAGGTG |
| fhuG_dw_seq | ACTCTGGGTGCGCAATTAAC |
| spm_seq1 | CATCAAAATACATCAATACACCTTCA |
| spm_seq2 | CCAGAATCAATAGAAATACGAAAAA |
| pchF_seq_up | CGGATCGCCCTGGTC |
| pchF_seq_dwn | GTCCAGGGTGGCGAAAC |
| pchcl_seq_up | ACAGGAGCGCACCGAAT |
| pchcl_seq_dwn | TCGTCGAGGCTGAACAGATT |
| pvdI_up_seq1 | GACGATCAGGACGAAGAACC |
| pvdI_dwn_seq1 | CAATCGCTGTTCGTCGAGT |
| **plasmid primer** | |
| pGPI_F | AGCTGATCCGGTGGATGAC |
| pGPI_R | ACGGTTGTGGACAACAAGC |

Restriction sites are underlined

**Table S3:** solvents used for solvent screen

| | |
|---|---|
| solvent 1 | dimethyl formamide |
| solvent 2 | dichloromethane |
| solvent 3 | ethyl acetate |
| solvent 4 | ethyl acetate + 0.1 % formic acid |
| solvent 5 | methanol |
| solvent 6 | water |
| solvent 7 | acetonitrile + water (1:1) |
| solvent 8 | acetonitrile + water (1:1) + 0.1% formic acid |
| solvent 9 | dimethyl formamide + water |
| solvent 10 | dimethyl formamide + water + 0.1 % formic acid |
| solvent 11 | Methanol + water (1:1) + 0.1% formic acid |
| solvent 12 | ethyl acetate + methanol (1:1) |
| solvent 13 | ethyl acetate + methanol (1:1) + 0.1 % formic acid |
| solvent 14 | n-Butanol |
| solvent 15 | n-Butanol + 0.1% formic acid |
